# CViewer: A Java-based statistical framework for integration of shotgun metagenomics with other omics datasets

**DOI:** 10.1101/2023.06.07.544017

**Authors:** Orges Koci, Richard K. Russell, M. Guftar Shaikh, Christine Edwards, Konstantinos Gerasimidis, Umer Zeeshan Ijaz

## Abstract

We have developed CViewer, a java-based framework to consolidate, visualize, and explore enormous amount of information recovered from shotgun sequencing experiments. This information includes and integrates all levels of gene products, mRNA, protein, metabolites, as well as their interactions in a single platform. The software provides a single platform to give statistical inference, and employs algorithms, some borrowed from numerical ecology literature to allow exploratory as well as hypothesis driven analyses. The end product is a highly interactive toolkit with multiple document interface, that makes it easier for a person without specialized knowledge to perform analysis of multiomics datasets and unravel biologically relevant hypotheses. As a proof-of-concept, we have used CViewer to explore two distinct metagenomics datasets: a dietary intervention study to understand Crohn’s disease changes during a dietary treatment to include remission, as well as a gut microbiome profile for an obesity dataset comparing subjects who suffer from obesity of different aetiologies and against controls who were lean.

## Introduction

While advances in high-throughput sequencing technologies have revolutionized the study of uncultured microbial communities, recent microbiome surveys using whole-genome shotgun sequencing delineates an enormous amount of information about microbial genomes, their taxonomic and functional profiles. With the ability to assemble nearly-complete genomes, we are now at the stage where, we can understand the mechanistic underpinnings to the biological processes involved in the studies for which the sequencing data are generated. Additionally, we can use metadata (typically, physicochemical and clinical parameters) relevant to the study to highlight patterns of interest. Whilst, many tools have become available in the past few years, from microbial genome binnings^1–3^, to annotation^4,5^, as well as downstream statistical analyses^6,7^, there is still a dire need to consolidate all the biological entities of microbial genomes, such as gene products, mRNA, protein, metabolites as well as their interactions in a single platform, and to enable exploration of this multi-component dataset in an easy-to-use interface.

This combination multiomics approach will be advantageous and will enable researchers, especially those who have a basic understanding about microbiology and microbial informatics, to quickly elucidate the interplay between microbiomes and their environments. Furthermore, combining visualizations with statistical inference in a single software platform has an added advantage of leveraging time spent on the analyses. Previous work towards this goal included conventional genome viewers like MGAViewer^8^ where, it is only possible for one to analyse at most two genomes together for comparative genomics, *albeit*, impractical for metagenomics exploration. Whilst there are other attempts such as Anvi’o^3^, a tool that implements an advanced analysis and visualization platform for omics data, combining taxonomic coverages with phylogenetic trees, and is typically used as a *de facto* tool for metagenomics exploration, as well as Elviz^9^ and WHAM!^10^ which, allow the exploration of metagenomic data. Although useful, these tools have limited support for statistical analyses, particularly Anvi’o^3^ and Elviz^9^, without any means to do integrative assessment. In view of the limitations as above, there is an unmet need for improvement to fully exploit the potential of the shotgun metagenomics data, both in terms of graphical user interface interactivity, as well as statistical inference. This serves as the basis for developing CViewer in a language such as Java that is not only cross-platform compatible, but also has established graphical interface to show relevant information on the fly.

The design principles of CViewer, include Java-based solution able to run on users’ local machines to incorporate the output from CONCOCT^2^, a binning software, as well as annotation data for the contigs from major third party taxonomic and annotation tools^5,11^. All this is done to explore the sample space in the context of clinical metadata, with the interface given in Figure 1. Beyond visualizing contigs from CONCOCT software, and looking at how they cluster at species level, we have implemented a variety of statistical algorithms from numerical ecology literature, what is conventionally available in R’s vegan package^7^, to allow exploratory as well as hypothesis driven analyses, emphasizing functional traits of microbial communities and their phylogenetic signal to assess community assembly. The implemented functionalities in CViewer include: alpha (based on the Simpson Index^12^, Shannon Entropy^13^ and Pielou’s Evenness^14^) and beta diversity (Principal Component Analysis^15^, Multi-dimensional Scaling^16^) estimates; statistical procedures to test covariates in terms of correlation with microbial community structure (Fuzzy Set Ordination^17^) and the variability they account for (PERMANOVA^18^) estimates (Figure 1A); differential abundance (based on the non-parametric Kruskall-Wallis^19^ test) and correlation analyses (including the Pearson’s^20^, Kendall’s^21^ and Spearman’s^22^ coefficients) (Figure 1B); enrichment analyses of KEGG^23^ metabolic pathways (Figure 1C); phylogenetic alpha diversity indices (based on the Net Relatedness Index and Nearest Taxon Index^24^) (Figure 1D); visualization and analysis of coverage, clustering and functional diversity of metagenomics contigs (Figure 1E); and integrative techniques for the analysis of multi-omics datasets (including the DISCO-SCA^25^, JAVA^26^ and O2PLS^27^ algorithms) (Figure 1F), with details given in the Supplementary Note 1. The software and the accompanying data, along with video demos exploring these features are available at https://github.com/KociOrges/cviewer.

**Figure 1.**
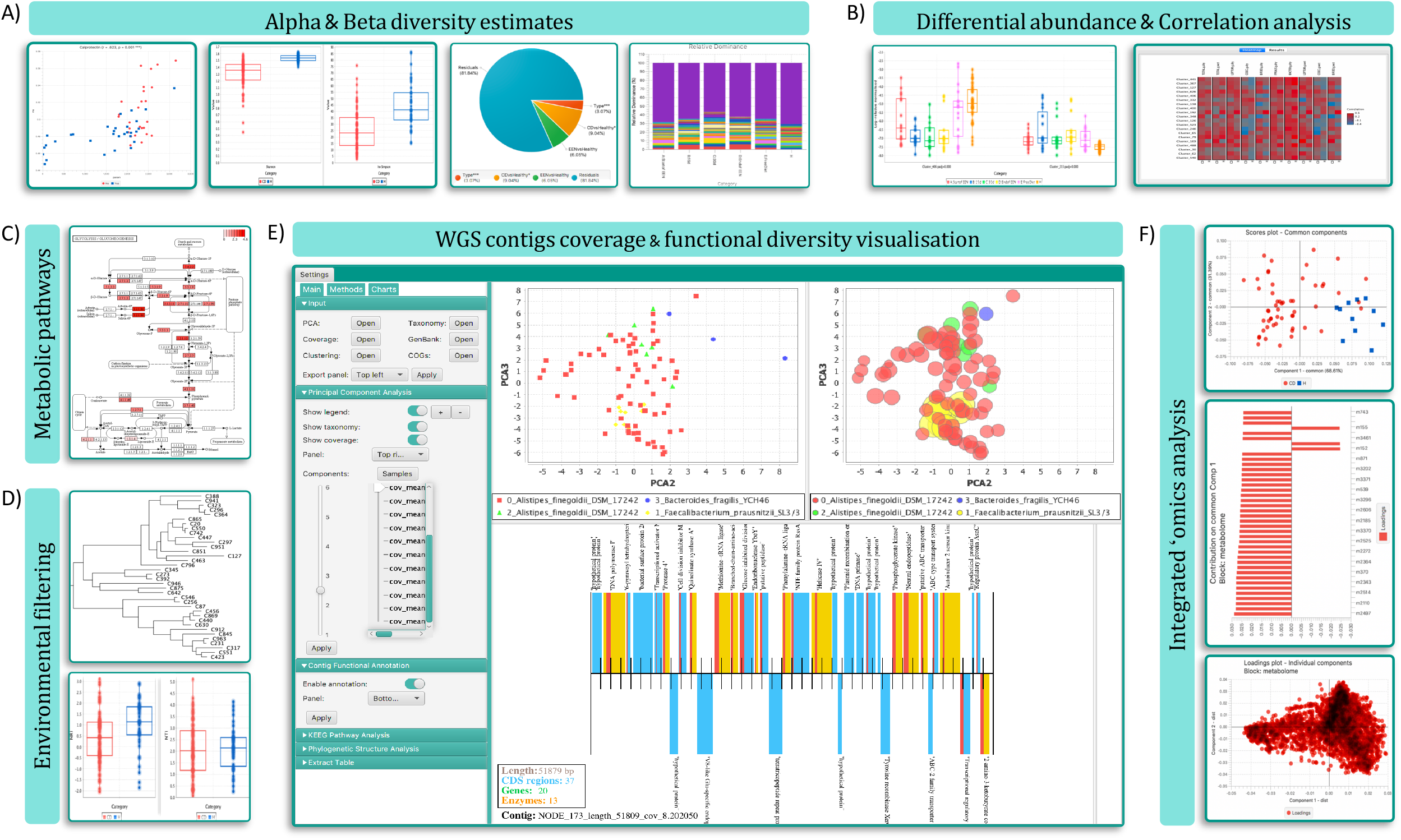
Schematic demonstrating the software layout and the main features that are supported in the system.

To demonstrate how CViewer is useful in analysing the metagenomics datasets, we have next used the software to explore a dataset of gut microbiome composition and metabolic profile from children with active Crohn’s disease (CD) who undergo dietary therapy with exclusive enteral nutrition (EEN). The data were categorized as: Crohn’s disease individuals who were treated with EEN for two months and healthy individuals to use as the reference microbiome and metabolome of healthy status. A smaller subset of these data has been previously published under Gerasimidis et al. 2014^28^, Quince et al. 2015^29^ and Alghamdi et al. 2018^30^, and additional samples and metagenomic sequencing were incorporated to see if the trends became more prominent. In addition, CViewer was tested on another dataset where the gut microbiome of children with pathological cause of hypothalamic obesity (Prader Willi Syndrome) was compared against children who suffer from common or classical type of obesity. Lean subjects from each of these two groups were used as controls. For more details on the datasets and the study characteristics the reader is referred to Supplementary Materials and Methods.

## Results

### Crohn’s disease dataset

#### Community structure of the gut metagenome and metabolome

Alpha diversity analysis using Shannon index suggested significantly lower diversity in the gut metagenome of CD children compared to that of healthy controls (p<0.0001) (Fig. 2A). Similar results were also found when we looked for differences during treatment with EEN (Fig. 2B & Supplementary Note 2). Shannon diversity remained significantly lower in CD subjects than in controls prior to EEN, throughout treatment and post-treatment (A:Start of EEN; p=0.0001, B:15d; p<0.0001, C:30d; p<0.0001, D:End of EEN; p<0.0001, E:Free Diet; p=0.0034). However, during EEN, the diversity of CD subjects decreased further, with the effect becoming noticeable after 15 days on treatment, reaching minimum diversity after ∼30 days, and then showing a slight recovery towards the end of EEN, and complete recovery to pre-treatment levels when patients returned to habitual diet. When we explored for differences between the treatment days, our results suggested a significant difference only between the timepoint C and E of EEN for Shannon’s diversity (p=0.0134).

**Figure 2.**
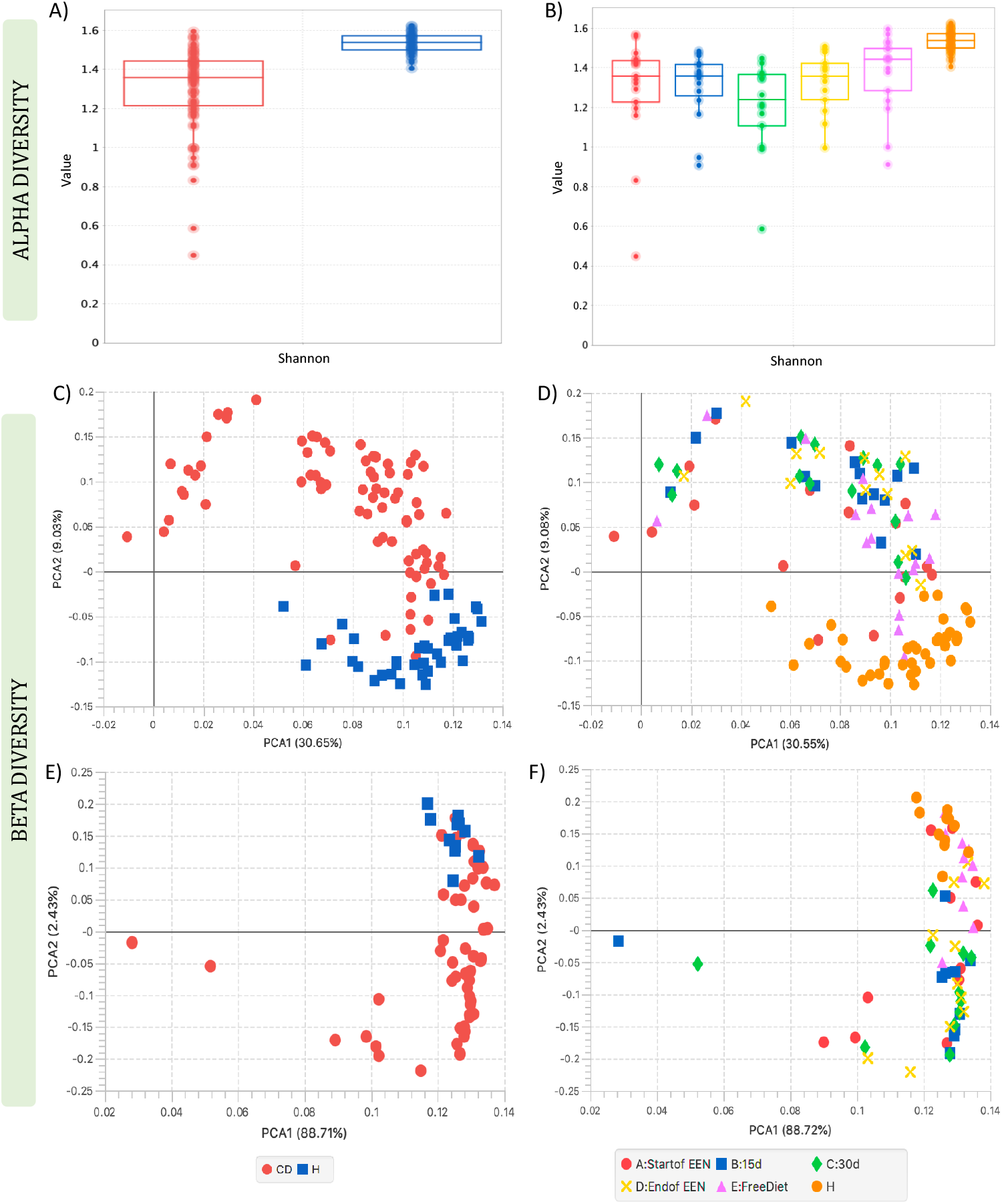
Figure demonstrating Shannon diversity of A) CD patients and healthy controls and B) of CD patients during EEN and healthy controls; PCA analysis of shotgun metagenomics community structure of C) CD patients and healthy controls, D) CD groups during EEN and healthy controls; and PCA analysis of the faecal metabolomic data derived from E) CD patients and healthy controls, and F) CD patients during EEN and healthy controls.

Beta diversity analysis of the gut metagenome using PCA showed an evident clustering of CD patients distinct from controls (Fig. 2C). This was also observed using PERMANOVA (distances between groups) which suggested that 8.3% of the variation in community structure was explained significantly by the sample groups (p = 0.001). When the CD subjects were grouped according to EEN samples, PCA described higher variability of the community structure for CD children compared to healthy controls (Fig. 2D) and the amount of significantly explained variability according to PERMANOVA increased to 11.22% (p = 0.001) suggesting that more differences in the community structure were accounted to changes during EEN. However, after the CD patients completed the treatment and returned to their free habitual diet, the samples appeared closer to the pre-treatment groups. Furthermore, analysis using Fuzzy Set Ordination (FSO) suggested a significant association between the microbial community structure of the CD individuals at point D (end of EEN) and E (free diet) of EEN with calprotectin (r=0.638, p=0.001), a marker of gut inflammation, (Supplementary Note 2 & Figs. S4A-S4B) and showed that patients who achieved remission, especially at the end of EEN, were related to lower calprotectin levels compared with the patients who still have active disease following completion of treatment. Similarly, PCA analysis on the faecal metabolome showed a distinct separation for the healthy control samples from the CD patient group (Fig. 2E). In addition, PCA showed that the metabolomes from healthy children were more tightly clustered, while CD patients were more variable and changed during EEN treatment (Fig. 2F).

#### Differential abundance analysis

Abundance analysis showed several clusters (genomes expressed in average abundances across samples) differentiating significantly in the CD groups and the healthy controls (Data_Table_ S1.xlsx). Among them, *Bifidobacterium longum* was in lower abundance in CD patients, at all sampling points, with a significant decrease noticed during EEN, when compared with the healthy control group (A:Start of EEN; p>0.05, B:15d; p=0.0002, C:30d; p=0.0001, D:End of EEN; p<0.0001) (Fig. 3A). Moreover, *Bifidobacterium adolescentis* was in significantly lower abundance in the CD samples at all sampling points with the effect being more evident after 15 days of EEN and onwards (A:Start of EEN; p=0.0282, B:15d; p=0.0002, C:30d; p=0.0002, D:End of EEN; p<0.0001, E:Free Diet; p=0.0002) (Fig. 3A). In a similar way, *Eubacterium rectale* was found in a lower abundance in CD groups and remained significantly lower at all points, although it moved closer to the control levels post EEN treatment on food reintroduction (A:Start of EEN; p=0.0001, B:15d; p<0.0001, C:30d; p<0.0001, D:End of EEN; p<0.0001, E:Free Diet; p=0.0357) (Fig. 3A). In contrast, *Escherichia coli* was significantly more prevalent in samples from CD patients at EEN initiation compared with the healthy controls (p=0.0009) and remained more abundant in CD individuals across all stages of EEN, although a decreasing pattern could be observed during the treatment course (Fig. 3A).

**Figure 3.**
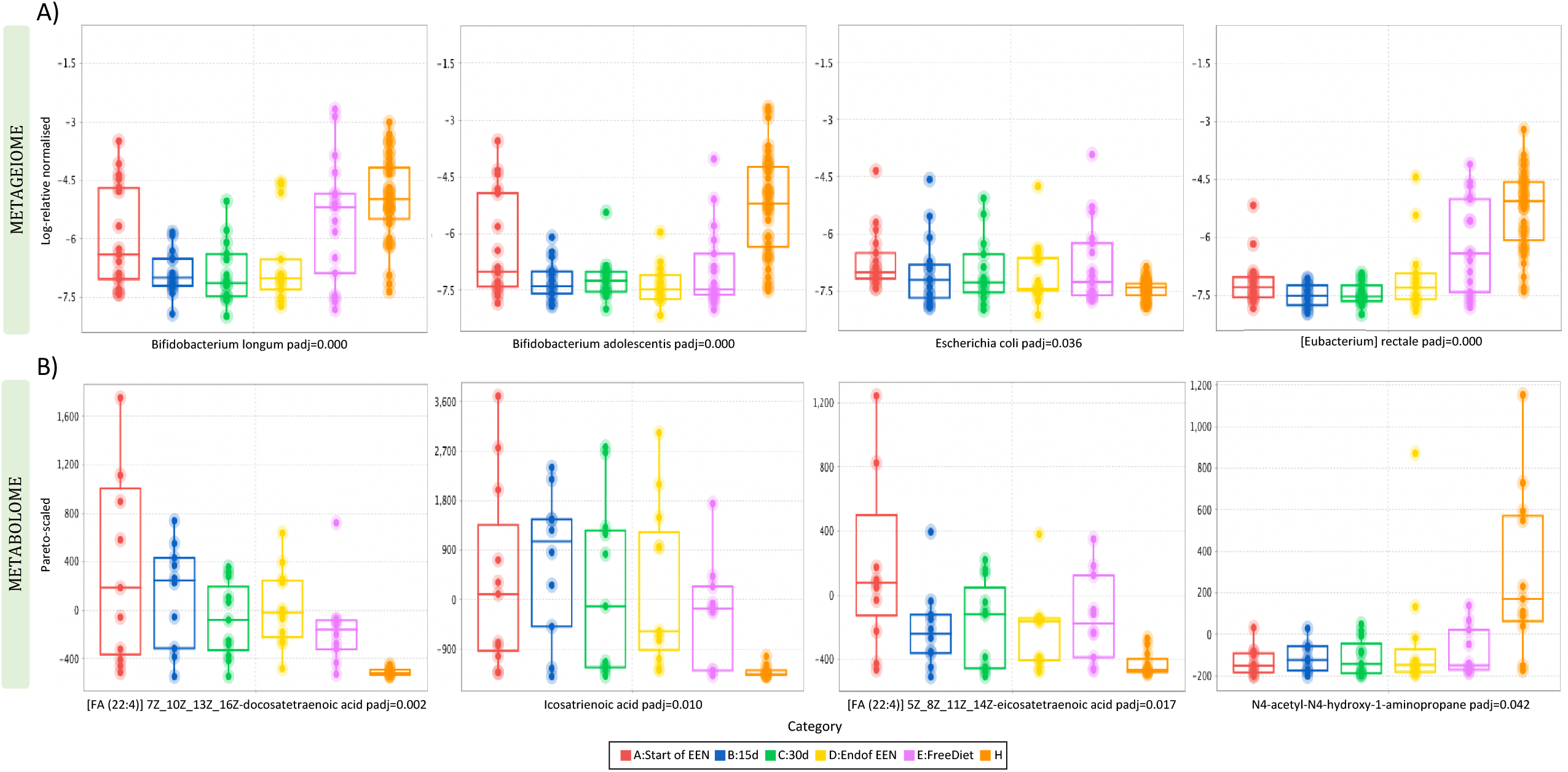
Figure showing sequential changes in A) the log-relative abundances of species and B) the pareto-scaled frequencies of metabolites that were significantly different in the CD groups (before, during, and after EEN treatment) and the healthy controls. Statistical significance is reported using corrected P-values (padj).

Differential analysis using Kruskal-Wallis also suggested that 487 annotated metabolites were significantly different between the CD groups and the healthy controls (Data_Table_S2.xlsx). More specifically, the levels of *docosatetraenoic acid* were found in a higher abundance in children with active CD at all sampling points compared to the healthy group (A:Start of EEN; p=0.0001, B:15d; p=0.0001, C:30d; p=0.0017, D:End of EEN; p=0.0002, E:Free Diet; p=0.0089). (Fig. 3B). This was also the case for *icosatrienoic acid* (A:Start of EEN; p=0.0014, B:15d; p=0.0002, C:30d; p=0.0026, D:End of EEN; p=0.0007, E:Free Diet; p=0.0147) and *arachidonic acid* (5Z_8Z_11Z_14Z-eicosatetraenoic acid) (A:Start of EEN; p=0.0001, B:15d; p=0.021, C:30d; p=0.0195, D:End of EEN; p=0.0189, E:Free Diet; p=0.0056) (Fig. 3B). The compounds remained significantly higher than the controls during EEN although some of the metabolites regressed to pre-treatment levels when subjects returned to free diet, with the effect being most pronounced for *arachidonic acid*. In contrast, the levels of an ornithine isomer (N4-acetyl-N4-hydroxy-1-aminopropane) were significantly lower in the CD samples than the healthy group, with the effect being more noticeable between the pre-treatment samples and the healthy controls (A:Start of EEN; p=0.0013, B:15d; p=0.016, C:30d; p=0.0021, D:End of EEN; p=0.0069, E:Free Diet; p=0.020) (Fig. 3B).

#### Changes in faecal bacterial metabolites during EEN

Significant differences were observed when we assessed the quantitative changes in the concentration of major targeted bacterial metabolites of CD patients during EEN (Fig. 4A). Compared to healthy individuals, faecal *pH* was significantly higher when patients were on EEN, but no differences were observed at EEN initiation or when patients returned to their habitual diet (A:Start of EEN; p>0.05, B:15d; p=0.0005, C:30d; p=0.0004, D:End of EEN; p<0.0001, E:Free Diet; p>0.5) (Fig. 4A). This was also the case for *total sulfide* (A:Start of EEN; p>0.05, B:15d; p=0.0015, C:30d; p=0.0004, D:End of EEN; p=0.0018, E:Free Diet; p>0.5) (Fig. 4A). In contrast, *butyric acid* was significantly lower in patients with CD compared to healthy controls with the effect being more evident after 15 days of EEN and onwards. This effect was lost when patients returned to their habitual diet (A:Start of EEN; p=0.0137, B:15d; p=0.0002, C:30d; p=0.0003, D:End of EEN; p<0.0001, E:Free Diet; p>0.5) (Fig. 4A). A similar pattern was also found for the proportional ratio of *butyric acid* (A:Start of EEN; p=0.0367, B:15d; p=0.0002, C:30d; p=0.0003, D:End of EEN; p=0.0002, E:Free Diet; p>0.5) (Fig. 4A).

**Figure 4.**
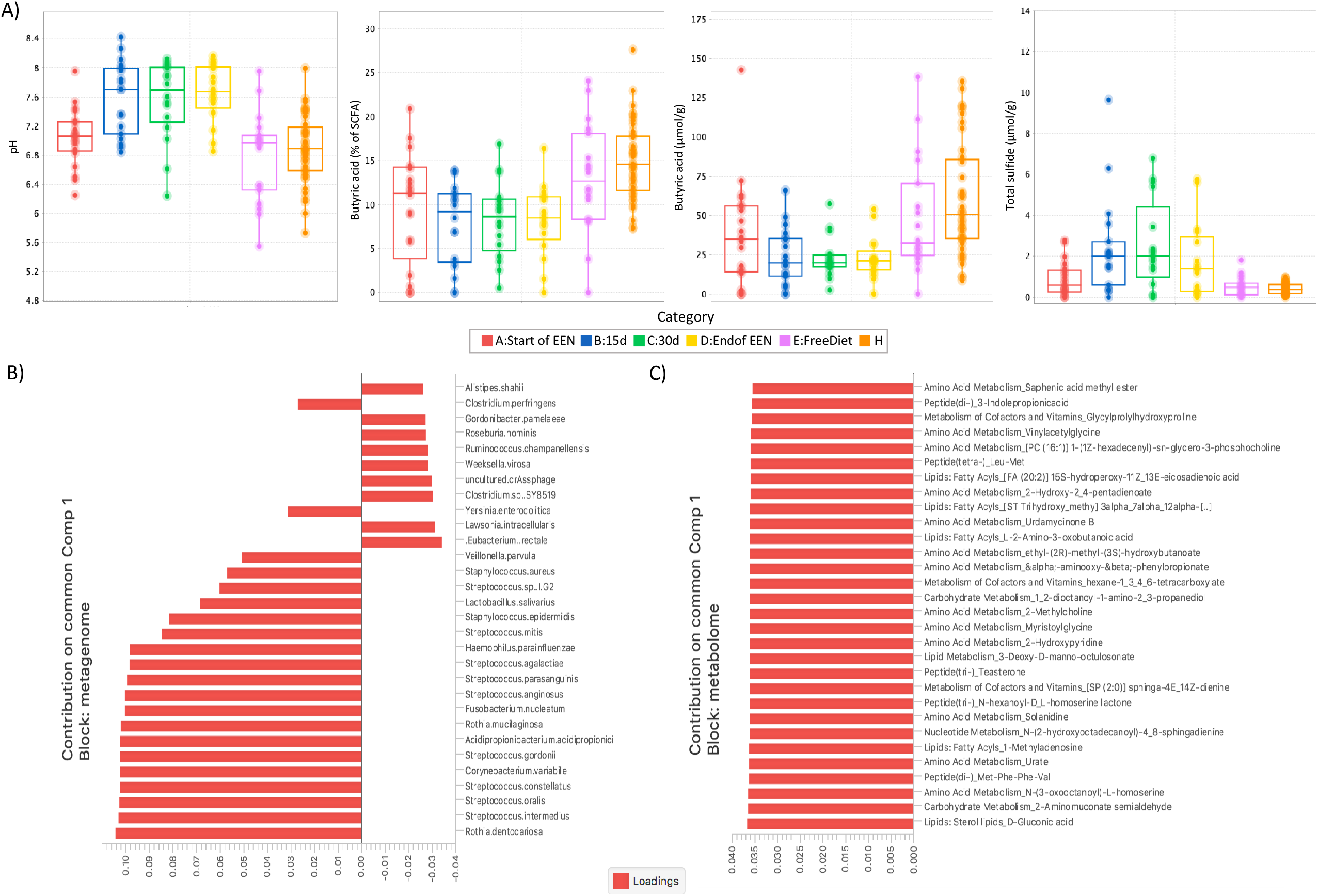
Figure demonstrating A) serial changes in faecal pH and concentration of major bacterial metabolites in CD subjects during EEN and healthy controls, B) the joint loadings for the top 30 species and C) top 30 metabolites with the highest contribution in the first component of the common structure. Loadings are sorted in decreasing order based on absolute values.

#### Integrated analysis of metagenomics and metabolomics

Integrated analysis of faecal metagenomics and metabolomics was performed to investigate for possible interactions between the two datasets. This methodology is particularly useful for decomposing the variability of the composite omics systems into a joint variability or common structure, that highlights the biological mechanisms underlying both the omics types under study. High contributions to the common structure across the two datasets (expressed as component loadings) could indicate a mechanistic association between the microbes and the metabolites, such as that a species may release a particular metabolite, or that specific metabolites may stimulate the growth of a particular species.

To explore this, the DISCO-SCA^25^ method was used for samples collected from CD patients before EEN initiation and healthy individuals (see Supplementary Material and Methods and Supplementary Note 1 for more details on methods and data pre-processing). Figures 4B and 4C visualize the estimated DISCO-SCA joint loadings for the omics datasets for the first common component, sorted by absolute value. When we explored the 30 first species with the highest contribution values for the metagenome, we noticed that *Rothia dentocariosa* had the largest value in the common structure loadings, while a high contribution was also noticed for several species of the genus *Streptococcus* (*Streptococcus intermedius/ oralis/ constellatus/ gordonii/ arginosus/ parasanguinis/ agalactiae*) (Fig. 4B). In a similar way, when we examined the 30 first metabolites with the highest contribution in the common structure for the metabolome, we found that *D-gluconic acid, 2-Aminomuconate semialdehyde* and *N-(3-oxooctanoyl)-L-homoserine* had the highest magnitude of positive loading values as compared to the other metabolites (Fig. 4C).

### Obesity dataset

#### Bacterial community structure

Shannon’s entropy and Pielou’s evenness showed no significant difference for the microbial diversity between the four groups, suggesting that the bacterial communities of the participants were similar in species abundance and richness (p=0.223) (Fig. 5A), and evenness (p=0.228). Shannon entropy provided similar results when participants were grouped according to obesity status (p=0.272) and obesity aetiology (p=0.551). Moreover, although the difference was not significant (p=0.156), it could be seen that the microbiota of the hypothalamic lean group was more diverse than the hypothalamic obese children (Fig. 5B). Beta diversity analysis using MDS showed no evident clustering in the community structure of the four groups, suggesting a similar microbial profile between the study participants (Fig. 5C). This was also confirmed by PERMANOVA, which suggested that the groupings did not account for a significant amount of the variability in the community structure (R2=7.13%, p=0.355). Using PERMANOVA, no significant effect of obesity phenotype (R2=1.66%, p=0.826) or obesity aetiology (R2=2.12%, p=0.532) on the community structure variation was observed. Moreover, differential analysis using Kruskal-Wallis did not suggest any significant differences in the average abundances of clusters (genomes) between the four groups when they were compared to each other, or when they were categorized according to obesity phenotype and aetiology.

**Figure 5.**
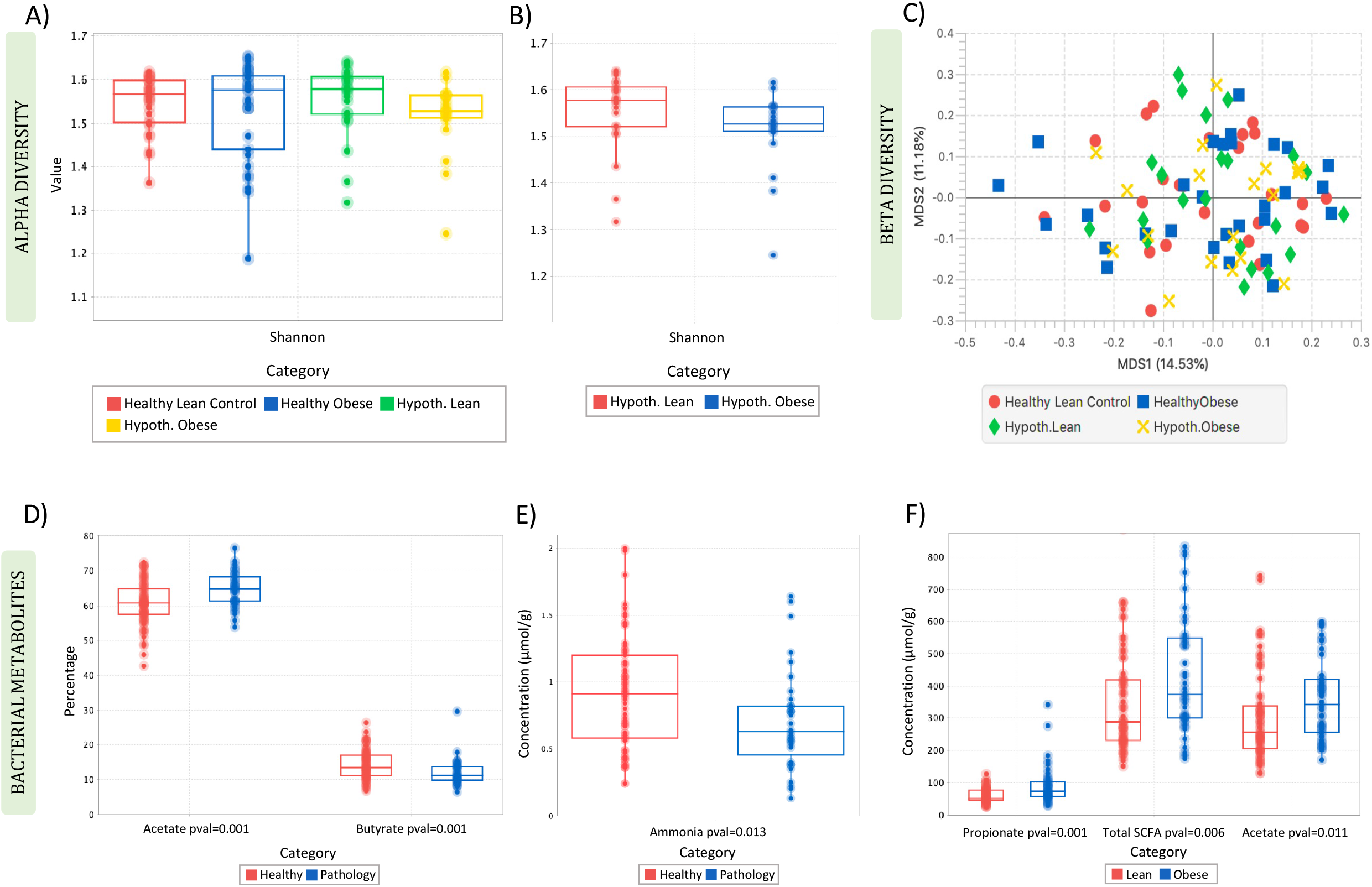
Shannon diversity of A) the 4 study groups, B) of hypothalamic lean (red) and hypothalamic obese participants (blue). Multidimensional scaling based on Bray-Curtis distance demonstrating clustering of the four study groups (C). Boxplots showing significant differences in the proportion of *acetate* (D) and concentration of *ammonia* (E) according to pathology and the concentration of major bacterial metabolites according to lean and obese phenotype (F). Lean; healthy and hypothalamic lean, obese; healthy and hypothalamic obese, healthy; healthy lean and obese, pathology; hypothalamic lean and obese.

#### Faecal bacterial metabolites

Significant differences in the proportion and concentration of bacterial metabolites were observed when the samples were grouped based on pathology (healthy vs. pathology). Healthy participants (healthy lean and healthy obese) had a significantly lower percentage (p=0.001) of *acetate* compared with the pathology groups (hypothalamic lean and hypothalamic obese) (Fig. 5H). In contrast, healthy participants had a significantly higher percentage of *butyrate* (p=0.001) and *ammonia* concentration (p=0.013) than the hypothalamic individuals (Figs. 5H-5I). In a similar way, when we examined for differences between the samples according to obesity status (healthy lean and hypothalamic lean vs. healthy obese and hypothalamic obese), it was noticed that the obese group (healthy obese and hypothalamic obese) had a significantly higher concentration (p=0.011) of *acetate*, concentration (p=0.001) and percentage (p=0.001) of *propionate* and concentration (p=0.006) of total *SCFA* than the lean samples (Fig. 5I & Fig. S5).

## Discussion

CViewer has successfully revealed patterns of interest in two independent datasets: a dataset for longitudinal gut microbiome and metabolomic profiles from children with Crohn’s disease who undergo dietary treatment with EEN; as well as on gut microbiome profile for an obesity dataset where subjects have a non-pathological or a pathological cause of obesity, and as compared to those who are lean. In the former study, beta diversity analysis of the gut microbiome and metabolome showed a clear separation of the CD groups throughout treatment and post-treatment from the healthy controls. In addition, the gut microbiota of CD patients changed soon after EEN initiation, was less diverse than in controls at all sampling points and decreased during EEN with a recovery to pre-treatment levels when patients returned to their habitual diet. These observations align with previous studies^31–34^, including some of ours^29,30^, demonstrating a distinct clustering of the metabolomes^34^ and metagenomes of CD patients from that of healthy controls and exhibiting general decreases in microbiota diversity relative to healthy individuals^31^, especially during EEN treatment^32,33^, suggesting the usefulness of an integrative approach. Our analysis also showed that classic commensal organisms such as *Bifidobacterium longum, Bifidobacterium adolescentis* and *Eubacterium rectale* differentiated in the CD groups over the course of EEN and were in lower abundance in CD children, compared to controls. These results align to previous research exploring the role of these organisms in CD pathogenesis^35^. Conversely, we noticed a higher abundance of *E. coli* in CD subjects than in controls; an expected outcome as the *E. coli* population is a much-studied topic in CD, particularly adherent invasive *E. coli* which are overrepresented in CD patients^36^.

Also, in accordance with our previous findings^30^, omega-6 fatty acids, including *docosatetraenoic acid* and *arachidonic acid*, were in significantly higher levels in the CD patients compared to healthy individuals and remained high both pre- and post-EEN treatment in the CD group. These findings conform to published work that highlight these compounds as pro-inflammatory in the gut and implicated in IBD^37,38^. At the same time, by exploiting exploratory methods further provided from the software, such as FSO, we were able to highlight an association between calprotectin and the microbiome of CD patients, especially for those who achieved remission at point D (end of EEN) of treatment. In their research, Quince et al.^29^ reported several taxa which were different between the CD and controls and significantly correlated with calprotectin, with *Bifidobacterium* spp. having the strongest negative and *Atopobium* spp. the strongest positive association with calprotectin in multivariate regression analysis. They also suggested that EEN caused the reduction in the relative abundance of gut bacteria that were positively and negatively associated with calprotectin. Although our study is limited in this perspective due to the lack of an implementation for regression analysis in the current version of the software, such an enhancement is one of our immediate future plans and with our findings for calprotectin and microbiome showing promising results, additional analysis on this aspect will be pursued in a future study with CViewer.

Finally, integrated analysis of the metagenome and the metabolome showed that *D-gluconic acid, 2-Aminomuconate semialdehyde* and *N-(3-oxooctanoyl)-L-homoserine* had the highest magnitude of positive loading values as compared to the other metabolites, while a high contribution was observed for *Rothia dentocariosa* and several species of the genus *Streptococcus* (*Streptococcus intermedius/ oralis/ constellatus/ gordonii/ arginosus/ parasanguinis/ agalactiae*) in the common structure loadings, i.e. the biological mechanisms underlying both omics datasets. *Rothia* species are assumed to have important interactions with the host immune system that may promote inflammatory diseases, including Crohn’s disease. *Rothia dentocariosa* in particular, has been previously reported to increase the production of the inflammatory cytokine tumour necrosis factor-alpha (TNF-a), and thus might act as an intermediate factor for increased inflammation in the oral cavity^39^. Moreover, *Streptococci*, especially the *Streptococcus anginosus* (milleri) group, comprising of *S. intermedius, S. arginosus and S. constellatus* species, has been associated with liver abscess in patients with Crohn’s disease^40^. Similar to the metabolite loadings, the values for these species were also positive and could suggest a potentially synergistic positive interaction between the microbes and the metabolites, where the highly contributing metabolites could either promote the growth of the species highlighted above, or that those species may produce the particular metabolites. These associations captured with CViewer provide some novel insights into the interactions of the two omics datasets and as relevant studies are still not prevalent in literature, supplementary correlation analysis (using e.g. Spearman’s correlation) of the metabolites and the species with high contributions in the common structure loadings could be useful to illustrate them even further as part of a future exploration with the tool.

In the latter study of lean and obese children of obesity of different aetiology, we demonstrated that the gut microbiota and metabolic activity did not differentiate between obesity of different aetiology or between obese and lean phenotypes, suggesting that gut microbiota is not incriminated in the aetiology of obesity. Moreover, our results described higher faecal *SCFA* in the two obese groups (common and hypothalamic obese) compared with lean (healthy and hypothalamic lean). Even though the role of *SCFA* in obesity is still not fully understood, recent research has demonstrated that higher concentration of *SCFA* in faeces is associated with obesity and hypertension^41^. However, the absence of difference in SCFA concentration between individuals with obesity of different aetiology strongly suggests that the increased production of SCFA in these groups is a secondary effect of differences in dietary patterns and hyperphagia rather than a primary defect increasing risk of obesity onset.

## Conclusions

Analytics for shotgun sequencing metagenomics has seen substantial growth in the recent years by availability of numerous tools, that cover and advance a particular aspect of the analyses, with their usability confined to passive running of scripts that allow little or no interactivity. CViewer on the other hand allows one to explore the datasets and benefits from the interactivity offered by the graphical user interface, particularly implementation platform in Java is well suited towards this purpose. Secondly, we have incorporated comprehensive statistical tools in the framework to allow one place to process and analyse all the datasets without the need to use any third-party tools such as R or Matlab. Moreover, multiomics exploration allows seamless integration of metagenomics with other omics technologies, such as metabolomics which, reveal patterns that not only consolidate/correlate between multiple datasets but provide discriminatory cues for multiple treatment groups. This way, an over-arching linkage between multiple modalities fills in the gaps in our understanding of how microbes are behaving in the context of the hypothesis under which the data is generated. Whilst we are exploring community assembly using phylogenetic alpha diversity measures (NRI/NTI), some new methods have appeared in recent years, some that are extension of the above^42,43^, and if implemented, can give further insights into the community assemblage processes. Nonetheless, in the absence of these methods, the existing functionality is sufficient to reveal useful patterns in the metagenomics datasets. Moreover, whilst we have tested and demonstrated integration of metagenomics with metabolomics in CViewer, future versions can possibly extend the analysis to a simultaneous exploration of metagenomics with two or more modalities including proteomics and transcriptomics to enable far more extensive exploration of the data.

## Data availability

The code for CViewer software and the associated data are available at: https://github.com/KociOrges/cviewer

## Author’s Contributions

UZI and KG designed the study; UZI, KG, and RKR directed this study as supervisors for OK; OK wrote the software under the guidance of UZI and carried out the statistical analysis; OK and UZI wrote the manuscript; KG critically interpreted findings; RKR, CE, and MGT provided feedback on the manuscript and clinical relevance/translation of this work; All authors read, commented on, and approved the paper.

## Acknowledgement

We would like to thank and dedicate this paper in memory of Dr Jaffar Khan who performed sample collection and laboratory analysis of the obesity datasets. We would also like to thank Christopher Quince who assisted with the generation of contigs for Crohn’s disease dataset.

## Supplementary Material

### Supplementary_Information.pdf

Supplementary Materials and Methods, Notes 1-3, Table S1, and Figs. S1–5.

### Data_Table_S1.xlsx

List of CONCOCT clusters (genomes) that were significantly different in the CD groups (before, during, and after EEN treatment) compared with the group of healthy controls (H) (P-value < 0.05). Mean expression indicates the mean log-normalized abundance for each genome. A post hoc pairwise Dunn’s comparison indicating significant differences between the groups is shown on the right half. A; StartofEEN; B; 15d; C; 30d; D; EndofEEN; E; FreeDiet; H; healhy.

### Data_Table_S2.xlsx

List of metabolites that were significantly different in the CD groups (before, during, and after EEN treatment) compared with the group of healthy controls (H) (P-value < 0.05). Mean expression indicates the mean Pareto-scaled frequency for each metabolite. A post hoc pairwise Dunn’s comparison indicating significant differences between the groups is shown on the right half. A; StartofEEN; B; 15d; C; 30d; D; EndofEEN; E; FreeDiet; H; healhy.

### Data_Table_S3.xlsx

Table showing significant differences in subject characteristics between subjects who suffer from obesity of different aetiology and against controls who are lean (P-value < 0.05). SDS; Standard Deviation Scores; Ht; Height (cm), Wt; Weight (Kg), BMI; Body Mass Index (kg/m^2^).

