## Supplementary_Information.pdf for "CViewer: A Java-based statistical framework for integration of shotgun metagenomics with other omics datasets"

#### 1. Materials and Methods

##### 1.1 Crohn's Disease dataset

###### 1.1.1 WGS dataset preparation

Faecal samples were collected from 23 children with CD (males: n=13; 6.9–14.7 years) during an 8-week course on EEN with a polymeric casein-based liquid feed (Modulen; Nestlé UK Ltd, York, United Kingdom). A maximum of five serial samples were collected per patient. The first sample (A) was collected before or within 6 days of EEN initiation (89% collected within 4 days), two during EEN (B:~16 & C:~32 days), and one close to end of treatment (D:~54 days). A final sample (E) was collected when patients returned to habitual diet (E:~63 days after EEN). For the number of participants at timepoints A, B, C, D and E, the total number of samples were 5, 3, 3, 2, and 10, respectively. Two samples were collected, at least 2 months apart, from 21 healthy children (males: n=12; 4.6–16.9 years) with no known family history of inflammatory bowel disease as a control group. The shotgun metagenomics samples were prepared with the Nextera XT Prep Kit (Illumina, FC-131- 1096, UK), and Illumina dual-barcoding Nextera XT Index kit (Illumina, FC-131-1002, UK). Sequencing libraries were pooled in equimolar concentration and quantified with the KAPA SYBR® FAST qPCR Kit (Kapa biosystems, KK4824, UK) then, loaded onto both lanes of a rapid run flow cell at 10 pM concentration. Clusters were generated on-board a HiSeq 2500 (Illumina) instrument and sequencing performed using TruSeq Rapid SBS Kit reagents (Illumina, FC-402-4001, FC-402-4002). Sequencing was performed following a paired-end 150 cycle recipe.

The above data have been previously published under Quince et al.[1], where the gut microbiota of 69 shotgun metagenomics faecal samples was explored using metagenomics reads for species taxonomic profiling using MetaPhlAn software. In this study, we reanalysed this dataset using additional samples and investigated the microbiome of 117 faecal extracts collected from both disease and healthy individuals. Metagenomics reads were assembled and grouped into bins using CONCOCT pipeline, which resulted in 976 genomic bins from a total number of 636,062 assembled contigs. Taxonomic classification of the CONCOCT bins was performed next using Kraken software.

##### 1.1.2 Bacterial metabolites

Faecal calprotectin concentration, a proxy marker of colonic inflammation, was measured by ELISA (PhiCal, Lysaker, Norway)[2]. Faecal pH was measured in 1:3 w/v slurries and ammonia with an automated ammonia analyser (HI 93715; Hanna, Bedfordshire, United Kingdom). Faecal water was calculated after freeze-drying samples. Short-chain fatty acids (SCFA; C2-C8) and branched chain fatty acids (BCFA; iso-butyrate, iso-pentanoic, and iso-caproic) were measured by gas chromatography[3]. Free and total faecal sulphide were measured by a modified spectrophotometric method. The concentrations of sulphide in samples was calculated by reference to calibration curves[4]. All these details have been published under Gerasimidis et al. 2014[5].

##### 1.1.3 LC-MS analysis

Metabolomic profiling using high resolution mass spectrometry with hydrophilic interaction chromatography was applied to 11 faecal extracts from eleven healthy children and to 54 faecal extracts from eleven children undergoing exclusive enteral nutrition for the treatment of active Crohn's disease (CD) at timepoints before, during (15, 30, and 60 days), and after treatment. In total, 4,255 metabolites were identified using LC-MS. Mobile phase solvents were freshly prepared and stored at room temperature for up to 48 h. Mobile phase A: ammonium carbonate buffer (20 mM, pH 9.2) was prepared by the addition of 1.92 g of ammonium carbonate to 800 mL of HPLC-grade water, followed by an adjustment to pH 9.2 with ammonia solution and then filled to a volume of 1 L. Mobile phase B: HPLC-grade acetonitrile only. The metabolites were eluted from the ZICpHILIC column (150 × 4.6 mm, 5 μm particle size) supplied by Hichrom Ltd. (Reading, UK) with a mobile phase consisting of 20 mM ammonium carbonate in HPLC-grade water (solvent A) and acetonitrile (solvent B), at a flow rate of 0.3 mL/min. All these details have been previously published under Alghamdi et al. 2018[6].

##### 1.1.4 Statistical analysis

Statistical analysis was performed in CViewer software. Metagenomics analysis was performed on log-relative normalized data and beta diversity was assessed using the reduced-order representation of the datasets using principal component analysis, with an additional standardization step by centring and scaling the data before doing PCA. For the evaluation of alpha diversity, we have used the Shannon's and Pielou's diversity index. Analysis of the variance for explanatory variables (or sources of variation) was performed using PERMANOVA. Relationships between community composition and clinical factors were assessed using fuzzy set ordination (FSO). To identify features that were significantly different between the conditions, Kruskal-Wallis test was used. The Benjamini-Hochberg correction was used on the returned P-values to correct for multiple testing and Dunn's test as a post-hoc procedure for pair-wise comparisons, where appropriate. PCA analysis of the metabolome was performed on log-2 transformed data and differential analysis of metabolites was performed on Pareto scaled frequencies. Differential analysis using Kruskal-Wallis statistic was also performed on each metabolite having an associated identification confidence level >= 5. This resulted in 1018 metabolites out of the initial 4,255, out of which 487 were significantly different between the CD groups (before, during, and after EEN treatment) and the healthy controls (Data_Table_S2.xlsx).

##### 1.1.5 Integrated analysis of metagenomics and metabolomics

For the integrated analysis of the metagenome and the metabolome, the DISCO-SCA algorithm was used (see Supplementary Note 1 for more details). To investigate for features that discriminate the CD patients from the healthy controls and for possible associations between the two omics datasets, the samples before EEN initiation were considered for analysis. Uncharacterized clusters/metabolites were dropped and metabolites with a confidence level >= 5 were considered. Prior to downstream statistics, the data were pre-processed, and the blocks were scaled, centred and weighted to correct for differences in the scales of the variables and the size of the datasets. To identify the optimal number of common and individual components, the *Model Selection* procedure (see Supp. Note 1) was used which yielded a total number of 2 common components for both datasets, and 10 individual components for each block. This information was then provided as an input to the algorithm.

#### 1.2 Obesity dataset

##### 1.2.1 WGS dataset preparation

The WGS dataset (unpublished as of now) comprised of 96 samples (simple obese n=13, healthy lean n=13, hypothalamic obese n=10, hypothalamic lean n=12 each for recruitment and after 2-3 months). The shotgun metagenomics samples were prepared using the methodology described in section 3.1.2.1. Metagenomic assembly analysis was performed using CONCOCT which returned 529 clusters from a total number of 542,999 assembled contigs. Taxonomic classification of the CONCOCT clusters was performed using Kraken software.

##### 1.2.2 Subject characteristics

Two faecal samples (A: at the time of recruitment, B: after 2-3 months) with anthropometric, body composition, and 24 h dietary data were collected from each participant. Height, weight and body mass index (BMI) were expressed as standard deviation scores (SDS) as described in Gerasimidis et al.[7].

##### 1.2.3 Bacterial metabolites

Faecal short chain fatty acids, hydrogen sulphide (total, free, and bound form), lactate (D, L, and total), ammonia, and faecal pH were measured as described in previous section. Briefly SCFA were measured in diethyl ether extracts using gas chromatography with flame ionisation detector (GC-FID). Hydrogen sulphide (free, total, and bound) was measured with a colorimetric assay according to the methylene blue reaction, ammonia using automated ammonia analyser (HANNAH Electrical HI93715, UK) and lactate (D, L, and total isomers) with an enzymatic assay. Values for SCFAs and BCFA in freeze dried faecal material were expressed as µg/g dry faeces. Free, bound, and total sulphide were expressed as µmol/g of dry faeces. Faecal NH3 and lactate were expressed as mg/g dry faeces.

##### 1.2.4 Statistical analysis

Statistical analysis was performed in CViewer software. The data were log-transformed, and beta diversity was assessed using the reduced-order representation of the datasets using Multidimensional Scaling based on Bray-Curtis distance. Alpha diversity was quantified by Shannon diversity index, which relates both species richness and evenness. Analysis of the variance for explanatory variables (or sources of variation) was performed using PERMANOVA. To identify features that were significantly different between the conditions, Kruskal-Wallis test was used. The Benjamini-Hochberg correction was used on the returned P-values to correct for multiple testing and Dunn's test as a post-hoc procedure for pair-wise comparisons, where appropriate. In addition, for demonstration purposes and to increase power, the A (recruitment) and B (repeated samples after 2-3 months) samples were merged into one group in downstream statistics.

### 2. Supplementary Notes

#### Supplementary Note 1: Information on third-party tools and statistical algorithms utilized in CViewer

##### 2.1.1 The CONCOCT software

CONCOCT is a toll useful for binning contigs by coverage and composition. Compared to existing binning methods that make use only of the sequence composition (the '$k$-mer' content, or subsequences of length $k$, considered as species-specific signatures)[8], CONCOCT uses Gaussian mixture models to cluster the metagenomic contigs utilising both sequence composition and coverage across multiple samples. The number of clusters is automatically estimated based on a variational Bayesian approximation. The method starts with a filtering step where usually contigs only greater than 1,000 bp are considered. Each filtered contig $i = 1, \dots, N$ is then represented by a feature vector of dimension $M + V$, including coverage of dimension $M$ and k-mer composition of dimension $V$. The coverage indicates the average number of reads per base pair from each of $j = 1, \dots, M$ samples mapping to that contig. The composition vector represents the frequency of each $k$-*mer* and its reverse complement for the same contig. Following the notation in[9] let $Y_i = \left(Y_{i,1}, \dots, Y_{i,M}\right)$ denote the coverage vector and $Z_i = \left(Z_{i,1}, \dots, Z_{i,V}\right)$ the composition vector for each contig. Prior to normalization, nonzero entries are removed by adding a small pseudo-count to both coverage and composition vectors so that the data can be log-transformed. For the coverage, the method adds a small value, i.e. $Y'_{i,j} = Y_{i,j} + 100/L_i$, where $L_i$ is the contig length and for the composition, the methods adds a single count to each $k$-*mer* $Z'_{i,j} = Z_{i,j} + 1$.

The coverage matrix $Y$ is first normalized over samples to account for different read numbers from a sample

$$Y''_{i,j} = \frac{Y'_{i,j}}{\sum_{k=1}^{N} Y'_{k,j}} \qquad \text{(Eq. S1)}$$

followed next by a normalization over contigs to obtain coverage profiles $p$. This accounts for coverage variation within a genome

$$p_{i,j} = \frac{Y''_{i,j}}{\sum_{k=1}^{M} Y''_{i,k}} \quad \text{(Eq. S2)}$$

The matrix $Z$ is also normalized to account for different contig lengths and calculate composition profile $q$

$$q_{i,j} = \frac{Z'_{i,j}}{\sum_{k=1}^{V} Z'_{i,k}} \quad \text{(Eq. S3)}$$

The coverage $p$ and composition $q$ are the combined into the feature matrix $X = [p \;\; q]^T$ of dimension $M + V$. Finally, CONCOCT reduces dimensionality using principal-component analysis on the log-profiles of $X$, and applies a Gaussian mixture model with a modified Bayesian model selection to cluster contigs into bins.

#### Table S1: Complete list of KEGG[10,11] metabolic categories that can be explored in CViewer

| **Carbohydrate metabolism** | Energy metabolism | Lipid metabolism | Nucleotide metabolism | Amino acid metabolism | Metabolism of other amino acids | Glycan biosynthesis and metabolism | Terpenoids and polyketides | Cofactors and vitamins | Biosynthesis of other secondary metabolites | Xenobiotics biodegradation and metabolism |
|---|---|---|---|---|---|---|---|---|---|---|
| Glycolysis / Gluconeogenesis | Oxidative phosphorylation | Fatty acid biosynthesis | Purine metabolism | Alanine, aspartate and glutamate metabolism | beta-Alanine metabolism | N-Glycan biosynthesis | Terpenoid backbone biosynthesis | Thiamine metabolism | Phenylpropanoid biosynthesis | Benzoate degradation |
| Citrate cycle (TCA cycle) | Photosynthesis | Fatty acid elongation | Pyrimidine metabolism | Glycine, serine and threonine metabolism | Taurine and hypotaurine metabolism | Various types of N-glycan biosynthesis | Monoterpenoid biosynthesis | Riboflavin metabolism | Stilbenoid, diarylheptanoid and gingerol biosynthesis | Aminobenzoate degradation |
| Pentose phosphate pathway | Photosynthesis - antenna proteins | Fatty acid degradation | | Cysteine and methionine metabolism | Phosphonate and phosphinate metabolism | Mucin type O-Glycan biosynthesis | Sesquiterpenoid and triterpenoid biosynthesis | Vitamin B6 metabolism | Flavonoid biosynthesis | Fluorobenzoate degradation |
| Pentose and glucuronate interconversions | Carbon fixation in photosynthetic organisms | Synthesis and degradation of ketone bodies | | Valine, leucine and isoleucine degradation | Selenocompound metabolism | Other types of O-glycan biosynthesis | Diterpenoid biosynthesis | Nicotinate and nicotinamide metabolism | Flavone and flavonol biosynthesis | Chloroalkane and chloroalkene degradation |
| Fructose and mannose metabolism | Carbon fixation pathways in prokaryotes | Cutin, suberine and wax biosynthesis | | Valine, leucine and isoleucine biosynthesis | Cyanoamino acid metabolism | Glycosaminoglycan biosynthesis - chondroitin sulfate / dermatan sulfate | Carotenoid biosynthesis | Pantothenate and CoA biosynthesis | Anthocyanin biosynthesis | Chlorocyclohexane and chlorobenzene degradation |
| Galactose metabolism | Methane metabolism | Steroid biosynthesis | | Lysine biosynthesis | D-Glutamine and D-glutamate metabolism | Glycosaminoglycan biosynthesis - heparan sulfate / heparin | Brassinosteroid biosynthesis | Biotin metabolism | Isoflavonoid biosynthesis | Toluene degradation |
| Ascorbate and aldarate metabolism | Nitrogen metabolism | Primary bile acid biosynthesis | | Lysine degradation | D-Arginine and D-ornithine metabolism | Glycosaminoglycan biosynthesis - keratan sulfate | Insect hormone biosynthesis | Lipoic acid metabolism | Indole alkaloid biosynthesis | Xylene degradation |

| | | | | | | | | | |
|---|---|---|---|---|---|---|---|---|---|
| Starch and sucrose metabolism | Sulfur metabolism | Secondary bile acid biosynthesis | Arginine biosynthesis | D-Alanine metabolism | Glycosaminoglycan degradation | Zeatin biosynthesis | Folate biosynthesis | Indole diterpene alkaloid biosynthesis | Nitrotoluene degradation |
| Amino sugar and nucleotide sugar metabolism | | Steroid hormone biosynthesis | Arginine and proline metabolism | Glutathione metabolism | Glycosylphosphatidylinositol(GPI)-anchor biosynthesis | Limonene and pinene degradation | One carbon pool by folate | Isoquinoline alkaloid biosynthesis | Ethylbenzene degradation |
| Pyruvate metabolism | | Glycerolipid metabolism | Histidine metabolism | | Glycosphingolipid biosynthesis - lacto and neolacto series | Geraniol degradation | Retinol metabolism | Tropane, piperidine and pyridine alkaloid biosynthesis | Styrene degradation |
| Glyoxylate and dicarboxylate metabolism | | Glycerophospholipid metabolism | Tyrosine metabolism | | Glycosphingolipid biosynthesis - globo and isoglobo series | Type I polyketide structures | Porphyrin and chlorophyll metabolism | Acridone alkaloid biosynthesis | Atrazine degradation |
| Propanoate metabolism | | Ether lipid metabolism | Phenylalanine metabolism | | Glycosphingolipid biosynthesis - ganglio series | Biosynthesis of 12-, 14- and 16-membered macrolides | Ubiquinone and other terpenoid-quinone biosynthesis | Caffeine metabolism | Caprolactam degradation |
| Butanoate metabolism | | Sphingolipid metabolism | Tryptophan metabolism | | Lipopolysaccharide biosynthesis | Biosynthesis of ansamycins | | Betalain biosynthesis | DDT degradation |
| C5-Branched dibasic acid metabolism | | Arachidonic acid metabolism | Phenylalanine, tyrosine and tryptophan biosynthesis | | Peptidoglycan biosynthesis | Biosynthesis of type II polyketide backbone | | Glucosinolate biosynthesis | Bisphenol degradation |
| Inositol phosphate metabolism | | Linoleic acid metabolism | | | Other glycan degradation | Biosynthesis of type II polyketide products | | Benzoxazinoid biosynthesis | Dioxin degradation |
| | | alpha-Linolenic acid metabolism | | | | Tetracycline biosynthesis | | Penicillin and cephalosporin biosynthesis | Naphthalene degradation |

| | | | |
|---|---|---|---|
| Biosynthesis of unsaturated fatty acids | Polyketide sugar unit biosynthesis | Carbapenem biosynthesis | Polycyclic aromatic hydrocarbon degradation |
| | Nonribosomal peptide structures | Monobactam biosynthesis | Furfural degradation |
| | Biosynthesis of siderophore group nonribosomal peptides | Clavulanic acid biosynthesis | Steroid degradation |
| | Biosynthesis of vancomycin group antibiotics | Streptomycin biosynthesis | Metabolism of xenobiotics by cytochrome P450 |
| | | Neomycin, kanamycin and gentamicin biosynthesis | Drug metabolism - cytochrome P450 |
| | | Acarbose and validamycin biosynthesis | Drug metabolism - other enzymes |
| | | Puromycin biosynthesis | |
| | | Novobiocin biosynthesis | |
| | | Staurosporine biosynthesis | |
| | | Aflatoxin biosynthesis | |

##### 2.1.2 Alpha diversity indices

***Shannon's (1948) diversity (H')***

Assume a community dataset of $N$ individuals partitioned into $S$ categories (species) and denote by $N_i$ the quantity (abundance) of $i$-th individual $(i = 1, \dots, S)$, where $N = N_1 + \dots + N_S$. Moreover, let $p_i = N_i/N$ denote the relative quantity of the $i$ -th individual, where $p_1 + \dots + p_S = 1$. Now, for the Shannon index to be computed, we first need to estimate the ratio of individuals $i$ over the total number of individuals $(p_i)$. Next, the product $(\mathrm{p}_i \log_b p_i)$ is summed across all species:

$$H' = -\sum_{i=1}^{S} p_i \log_b p_i \tag{Eq. S4}$$

The natural log makes the summation negative, and thus, the product $(\mathrm{p}_i \log_b p_i)$ is multiplied by -1. The $H'$ value now accounts for both richness and evenness and increases as both these amounts increase.

***Simpson's (1949) diversity (D₁)***

The Simpson's index is obtained by calculating the ratio of individuals $i$ over the total number of individuals ( $p_i$ ), squaring and summing them across all the species

$$D = \sum_{i=1}^{S} p_i^2 \tag{Eq. S5}$$

Now as $D$ increases, diversity decreases. For this reason, since $D$ can have a value between zero and one, it is often preferred that it is described by its complement:

$$D_1 = 1 - \sum_{i=1}^{S} p_i^2 \tag{Eq. S6}$$

***Inverted Simpson's diversity ($D_2$)***

The inverse of Simpson's original index ($D_1$) also increases as diversity increases.

$$D_2 = \frac{1}{\sum_{i=1}^{S} p_i^2} \tag{Eq. S7}$$

***Pielou's (1969) Evenness***

The value for this index is calculated based on Shannon's diversity, as Shannon accounts for both richness and evenness of the community. The statistic can have a value between 0 and 1, with low evenness values indicating that the species in a microbial community are relatively less equal in abundance, and high values indicating that abundances of species are relatively similar. The index is then calculated as

$$J = \frac{H'}{\log(S)} \qquad \text{(Eq. S8)}$$

where $H'$ denotes Shannon's diversity.

##### 2.1.3 Correlation coefficients

***Pearson's correlation***

Given a pair of variables ($X$ and $Y$) the Pearson's correlation coefficient is calculated as

$$p_{XY} = \frac{Cov(X,Y)}{\sigma_X * \sigma_Y} \qquad \text{(Eq. S9)}$$

where $Cov(X,Y)$ is the covariance of $X$ and $Y$ defined as

$$Cov(X,Y) = \frac{1}{N-1} * \sum_{i=1}^{N}[(X_i - \bar{X})(Y_i - \bar{Y})] \qquad \text{(Eq. S10)}$$

$\sigma_X$ and $\sigma_Y$ denote the standard deviation for $X$ and $Y$, where $\sigma_X$ is defined as

$$\sigma_X = \sqrt{\frac{1}{N-1} * \sum_{i=1}^{N}(X_i - \bar{X})^2} \qquad \text{(Eq. S11)}$$

and $\sigma_Y$ is defined as

$$\sigma_Y = \sqrt{\frac{1}{N-1} * \sum_{i=1}^{N}(Y_i - \bar{Y})^2} \qquad \text{(Eq. S12)}$$

where $\{X_1, X_2, \dots, X_N\}$, $\{Y_1, Y_2, \dots, Y_N\}$ are the observed values in $X$ and $Y$, $\bar{X}$ and $\bar{Y}$ is the mean value of these observations in $X$ and $Y$. The statistic gives a value that can range from -1, indicating perfect negative association, to +1, indicating perfect positive association. A value of zero suggests no association.

***Kendall's Tau-b rank correlation***

Kendall's tau coefficient estimates the correlation between two variables based on the number of concordances and discordances between the observations of these variables. Thus, two observations ($X_i$, $Y_i$) and ($X_j$, $Y_j$) are assumed *concordant* if the ranks for both elements agree, i.e, if $X_i > X_j$ and $Y_i > Y_j$, or if $X_i < X_j$ and $Y_i < Y_j$. In a similar way they are assumed *discordant* if both $X_i > X_j$ and $Y_i < Y_j$, or if $X_i < X_j$ and $Y_i > Y_j$.

The Kendall Tau-b coefficient is then calculated as

$$\tau_B = \frac{n_c - n_d}{\sqrt{(n_0 - n_1)(n_0 - n_2)}} \quad \text{(Eq. S13)}$$

where $n_c$ denotes the number of concordant pairs, $n_d$ the number of discordant pairs, and $n_0$ is the total number of pairs that can be produced for a sample of size $N$ and is calculated as

$$n_0 = \binom{N}{2} = \frac{1}{2}N(N-1) \quad \text{(Eq. S14)}$$

Next, $n_1$ is defined as

$$n_1 = \sum_{i=1}^{N} \frac{t_i(t_i - 1)}{2} \quad \text{(Eq. S15)}$$

where $t_i$ is the number of tied values in the $i^{\text{th}}$ group of ties in $X$, and $n_2$ is calculated in a similar way as

$$n_2 = \sum_{j=1}^{N} \frac{u_j(u_j - 1)}{2} \quad \text{(Eq. S16)}$$

where $u_j$ is the number of tied values in the $j^{\text{th}}$ group of ties in $Y$. The value of this statistic also ranges between -1 and +1, with a zero-value suggesting no association and positive and negative values suggesting positive and negative associations respectively.

***Spearman's rank correlation***

The Spearman's coefficient $r_s$ is the non-parametric alternative to Pearson's coefficient and is calculated on the ranks of the data rather than the original numeric values. The interpretation of the statistic is similar to the other coefficients. The calculation of the Spearman's correlation $r_s$ is performed based on the formula below

$$r_s = 1 - \frac{6\sum_{i=1}^{N} d_i^2}{N(N^2 - 1)} \quad \text{(Eq. S17)}$$

where $N$ is the number of the observations and $d^2$ the squared difference between two ranks of each observation. In cases when two or more observations were assigned to the same rank (tied ranks), then the Spearman correlation becomes

$$r_s = p_{rg_x rg_y} = \frac{Cov(rg_x, rg_y)}{\sigma_{rg_x} * \sigma_{rg_y}} \qquad \text{(Eq. S18)}$$

where p denotes the Pearson correlation coefficient using the ranks of the observations, $Cov(rg_x, rg_y)$ is the covariance and $\sigma_{rg_x}, \sigma_{rg_y}$ are standard deviations of the rank observations.

##### 2.1.4 Beta diversity and ordination techniques

***Multidimensional Scaling***

Given a matrix of pairwise distances, Multidimensional scaling (MDS) aims to reconstruct a space of low dimension that preserves these distances. The procedure can be divided into three main steps described right below:

1) The first step starts with the computation of a distance matrix containing the pairwise distances between the $N$ elements of a dataset (Fig. S1), which is calculated based on a given distance index. CViewer provides the methods for some of the most common distance measures, i.e. Bray-Curtis, Jaccard and Euclidean distance to generate the dissimilarity matrix for the given dataset. If abundance data is available, then Bray-Curtis distance is often recommended, while Jaccard index is mostly applied for presence/absence data. If we consider $x_{i,j}$ and $x_{i,k}$ as the abundance or presence/absence of species (column) $i$ and sites (rows) $j$ and $k$, then the Euclidean distance between $x_i$ and $x_j$ is given as

$$d_{j,k} = \sqrt{\sum_i \left(x_{i,j} - x_{i,k}\right)^2} \qquad \text{(Eq. S19)}$$

Accordingly, the Bray-Curtis distance between $x_i$ and $x_j$ is given as

$$d_{j,k} = \frac{\sum_i \left|x_{i,j} - x_{i,k}\right|}{\sum_i \left(x_{i,j} - x_{i,k}\right)} \qquad \text{(Eq. S20)}$$

In a similar way, Jaccard is computed as shown below

$$d_{j,k} = 2 * \frac{B_{j,k}}{1 + B_{j,k}} \quad \text{(Eq. S21)}$$

where $B_{j,k}$, is Bray-Curtis dissimilarity.

2) Next, the distance matrix is squared and double-centred as described below

$$\widehat{D}_{i,j} = D_{i,j} - b_j - c_i$$
$$= D_{i,j} - \frac{1}{N}\sum_{k=1}^{N} D_{k,j} - \frac{1}{N}\sum_{k=1}^{N} D_{i,k} \quad \text{(Eq. S22)}$$

where $b$ denotes the row mean and $c$ the column mean accordingly. Next, the double centred matrix is computed using the squared distance matrix

$$C = -\frac{1}{2}\widehat{D} \quad \text{(Eq. S23)}$$

3) Then, the centred distance matrix is decomposed into the $k$ largest eigenvalues and eigenvectors of $C$, with $k$ denoting the number of the dimensions to be returned in the output. Finally, the classical MDS is computed as shown below

$$P = E_k\sqrt{\Lambda_k} \quad \text{(Eq. S24)}$$

where $\Lambda$ denotes the matrix of $k$ eigenvalues and $E_k$ the matrix of $k$ eigenvectors of $C$.

$$D = \begin{pmatrix} 0 & d_{1,2} & d_{1,3} & \cdots & d_{1,N} \\ d_{2,1} & 0 & d_{2,3} & \cdots & d_{2,N} \\ d_{3,1} & d_{3,2} & 0 & \cdots & d_{3,N} \\ \vdots & \vdots & \vdots & \ddots & \vdots \\ d_{N,1} & d_{N,2} & d_{N,3} & \cdots & 0 \end{pmatrix}$$

**Figure S1**: Given a set of $N$ elements (samples), the distance matrix $D$ is a $NxN$ square matrix containing the pairwise distances between these elements as calculated based on a given distance index (e.g. Euclidean, Bray-Curtis, Jaccard).

##### ***Fuzzy Set Ordination***

Fuzzy set ordination (FSO) expects that samples are assigned memberships in sets that can range from 0 to 1. For this reason, the method does not use the raw community data, but rather a similarity (or distance) matrix, which is calculated prior to the actual statistic. If two samples share no species, FSO assumes that a SI is 0, while if they have the same species in common, then SI is equal to 1. CViewer supports three different similarity indices namely, Baroni-Urbani

& Buser, Horn and Yule. Even though Baroni-Urbani & Buser and Yule are binary indices, they can be converted into an abundance SI[12]. The abundance SIs are described below. First, let us assume a community of $N$ individuals (samples) partitioned into $S$ categories (species) and indicate by $N_j$ the abundance of $j$-th category ($j = 1, \dots, S$), where $N = N_1 + \cdots + N_S$. Then the $Baroni - Urbani$ & $Buser$ similarity index is

$$\frac{A + C}{B + D} \qquad \text{(Eq. S25)}$$

where $A$, $B$, $C$ and $D$ are calculated as

$$A = \sum_{i=1}^{S} \min(x_{ij}, x_{ik}),$$

$$B = \sum_{i=1}^{S} \max(x_{ij}, x_{ik}),$$

$$C = \sqrt{\left(\sum_{i=1}^{S} min(x_{ij}, x_{ik})\right)\left(\sum_{i=1}^{S}\left(\max_j(x_{ij}) - \max(x_{ij}, x_{ik})\right)\right)},$$

$$D = \sqrt{\left(\sum_{i=1}^{S} min(x_{ij}, x_{ik})\right)\left(\sum_{i=1}^{S}\left(\max_j(x_{ij}) - \max(x_{ij}, x_{ik})\right)\right)},$$

Similarly, Horn index is calculated as

$$\frac{A - B - C}{D} \qquad \text{(Eq. S26)}$$

where $A$, $B$, $C$ and $D$ are defined as

$$A = \sum_{i=1}^{S}\left[(x_{ij} + x_{ik}) \log(x_{ij} + x_{ik})\right],$$

$$B = \sum_{i=1}^{S}(x_{ij} \log x_{ij}),$$

$$C = \sum_{i=1}^{S}(x_{ik} \log x_{ik}),$$

$$D = \left[(N_j + N_k) \log(N_j + N_k)\right] - N_j \log N_j - N_k \log N_k$$

And finally, Yule index is

$$\frac{A}{B + C} \qquad \text{(Eq. S27)}$$

where $A$, $B$ and $C$ are calculated as

$$A = \sqrt{\left(\sum_{i=1}^{S} min(x_{ij}, x_{ik})\right)\left(\sum_{i=1}^{S}\left(\max_j(x_{ij}) - \max(x_{ij}, x_{ik})\right)\right)},$$

$$B = \sqrt{\left(\sum_{i=1}^{S} min(x_{ij}, x_{ik})\right)\left(\sum_{i=1}^{S}\left(\max_j(x_{ij}) - \max(x_{ij}, x_{ik})\right)\right)},$$

$$C = \sqrt{\left(\sum_{i=1}^{S} max(x_{ij}, x_{ik}) - x_{ik}\right)\left(\sum_{i=1}^{S} max(x_{ij}, x_{ik}) - x_{ij}\right)}.$$

where for each index samples $j$ and $k$ are being compared, for species $i = 1$ to $S$, $x_{ij}$ is the quantity of species $i$ at sample $j$, $\max_j(x_{ij})$ = the maximum quantity of species $i$ across all samples $j$, and $N_j = \sum x_{ij}$. Even though these indices can provide us with an appropriate similarity matrix, FSO assumes that the data will be given as distances and thus, the final product is given as the complement of SI (1 - SI).

##### 2.1.5 Multiomics data integration

To this end, three different techniques have been proposed for a simultaneous analysis of multiple omics datasets, namely *Simultaneous Component Analysis with rotation to COmmon and DIstinctive components (DISCO-SCA)*[13], *Joint and Individual Variation Explained (JIVE)*[14] and *Two-way Orthogonal Partial Least Squares (O2PLS)*[15]. All of them are helpful for providing a "global" view on the biological system of interest by decomposing the variability of the composite omics datasets into a joint variability or common structure, that represents the mechanisms underlying all the omics datasets under study, and individual variability or distinctive structure, that represents the biological mechanisms underlying a single omics dataset. CViewer offers a JAVA implementation of the above methods providing support for an integrative analysis of multiomics datasets.

###### 2.1.5.1 Data pre-processing

In a simultaneous analysis of multiple datasets, it is expected that the different blocks comprising the data are linked with a common set of entities. This common set can either be the same set of *objects* or the same set of *variables*. More specifically, the data matrices are usually organized in such a way that the rows of the data correspond to the different objects describing the samples or experimental units (e.g., conditions) and the columns to the different variables (e.g., CONCOCT clusters/genomes, metabolites). In this work, the data of interest are organized in the same way, and they consist of two data matrices that have the same set of objects in common, i.e. the measurements from the different omics data were obtained for the same set of samples (e.g. Crohn's disease or Healthy individuals). Thus, the framework implemented in CViewer, supports an integrative analysis for two omics datasets that are linked by a common sample space.

Prior to analysis, it is useful that the data are pre-processed. As the data are generated from different omics technologies, they may be considerably different in size and describe features that are expressed in scales that are hard to compare, affecting this way the analysis results. For this reason, CViewer provides a number of pre-processing steps that can be applied to correct these differences. When the variables differ largely in scales (or abundance), one may consider centring the variables and/or scale them within each block to a sum of squares of one. In addition, weighting blocks together can be useful to avoid the effects of blocks having considerably different sizes.

##### 2.1.5.2 Model selection

Before ones chooses any of the integrated methods to perform component analysis, one must first provide the number of components that the dataset is expected to cover majority of the variation in this component space (such is the norm in algorithms similar to PCA which fit loading vectors and calculate scores on these loading vectors) along with their characterization as either common or distinctive. This information is required for each integrative approach (i.e., DISCO-SCA, JIVE and O2PLS) and it is necessary so that the common and individual structures can be successfully identified and separated from each other. For example, if we assume a component analysis for $K = 2$ data matrices $X_1, \dots, X_k$, $(k = 1, \dots, K)$ that is performed for $R = 3$ total components, then let $c$ denote the number of common components, and $d_k$ the individual components of the $k^{th}$ data block. Our model assumes the first component as common $(c = 1)$, the second as distinctive for the first data block $(d_1 = 1)$, and the third as distinctive for the second data block $(d_2 = 1)$. To estimate the number of common and distinctive components for the dataset of interest, CViewer provides the JAVA implementation for the *Model Selection* function as presented in the STATegRa R package[16] for multiomics data integration. Pre-processing of data should also be considered before the model selection analysis is performed (see previous section).

##### 2.1.5.3 Algorithms for multiomics data integration

To describe the algorithms for an integrative omics analysis, we will rely on the following notation: matrices and vectors are denoted by upper- and lower-case letters respectively. The objects $(i = 1 \dots I)$ represent the rows of the matrices $(I \times J)$ and the variables $(i = 1 \dots J)$ represent the columns. The transpose of the matrices and vectors is denoted by the superscript $^T$, and the cardinality of an index is indicated by the capital of the letter used to run the index. For example, if $k$ denotes the block, then the $k^{th}$ data block is indicated by $X_k$, with $k$ running from 1 to $K$. As CViewer supports an integrative analysis of two omics data types, the letter $K$ denoting the number of blocks is assumed to be 2 ($K = 2$) throughout this section.

For a joint analysis of $K = 2$ data matrices $X_1, \dots, X_k$, $(k = 1, \dots, K)$ each of dimension $(I \times J_k)$, DISCO-SCA, JIVE and O2PLS are based on the following decomposition for each $X_k$, for finding their common and distinctive variation

$$X_k = C_k + D_k + E_k, \quad k = 1, \dots, K \qquad \text{(Eq. S28)}$$

where $C_k$ $(I \times J_k)$ denotes the common part, $D_k$ $(I \times J_k)$ the distinctive part and $E_k$ $(I \times J_k)$ indicates the residual error. However, to accomplish this, each method follows a different approach by applying *Simultaneous Component Analysis* (SCA) and singular value decomposition (SVD). More details about the SCA and SVD methods are given below. SCA can be useful for resolving the biological mechanisms underlying the data, however, it does not define a method to separate common and distinctive components. To further identify joint and specific variation in a multi-block dataset, each of the integrative techniques follows a specific methodology which is described in the following sections.

*Simultaneous Component Analysis with rotation to COmmon and DIstinctive components (DISCO-SCA)*

Simultaneous component analysis does not explicitly separate common and distinctive variation but usually reflects a mix of common and distinctive information. To disentangle these kinds of information, Schouteden et al.[13] proposed the idea to reveal common and distinctive components for two different blocks of data, by a proper rotation of the components obtained from a simultaneous component analysis. For a given number of $R$ components, the method introduces an orthogonal rotation of the concatenated component loadings $P$ (common object mode) toward a common/distinctive structure, which is called *target matrix* (details given below). The matrix defines a mechanism that is distinctive for a single data block $X_k$, by block-specific loadings that are zero for all data blocks apart for those for which it is distinctive. The loadings for the common components are left undefined. Next, the loading matrix $P$ is rotated orthogonally towards the *target matrix* with optimal rotation matrix $B_{opt}(R \times R)$ which is calculated by minimizing the following function

$$B_{opt} \underset{min}{\longrightarrow} \sum\left(W \circ (PB)\right)^2 \quad \text{s.t } B^T B = \text{I} \qquad \text{(Eq. S29)}$$

with $W$ a weight matrix which contains ones for the positions that correspond to the zeros specified in the target matrix and zeros everywhere else, and where $\circ$ denotes the Hadamard product[1]. The rotated scores and loadings given in the output ($T_r = TB_{opt}$ and $P_r = PB_{opt}$) are then estimated using the $B_{opt}$ product. With this approach, only those parts targeted to be zero, i.e. the distinctive components, contribute to the optimization function, and the remaining parts have no influence on it. The final decomposition of the DISCO method for two data matrices $X_1$ and $X_2$ is

$$\begin{aligned} X_1 &= C_1 + D_1 + E_1 = T_c P_{c_1}^T + T_{d_1} P_{d_1}^T + E_1 \\ X_2 &= C_2 + D_2 + E_2 = T_c P_{c_2}^T + T_{d_2} P_{d_2}^T + E_2 \end{aligned} \quad \text{(Eq. S30)}$$

where $c$ and $d$ denote the components for the common and individual structure respectively. Note that the common scores $T_c$ are the same for both datasets.

*Target matrix for DISCO-SCA*

To illustrate the structure of a *target matrix* for an analysis with DISCO-SCA, consider a case with $K = 2$ data blocks and $R = 3$ components. We will assume that the first component is distinctive for the first data block ($d_1 = 1$), the second distinctive for the second data block ($d_2 = 1$), and the third one is common ($c = 1$). Then, the target matrix $P^{target}$ in our example, will be as follows

$$P^{target} = \begin{bmatrix} P_1^{target} \\ ------ \\ P_2^{target} \end{bmatrix} = \begin{bmatrix} \times & 0 & \times \\ \vdots & \vdots & \vdots \\ \times & 0 & \times \\ - & - & - \\ 0 & \times & \times \\ \vdots & \vdots & \vdots \\ 0 & \times & \times \end{bmatrix} \quad \text{(Eq. S31)}$$

where $P_1^{target}$, $P_2^{target}$ represent the loading matrices that refer to the two individual data blocks and $\times$ denotes an unspecified entry.

*Joint and Individual Variation Explained (JIVE)*

JIVE[14] is another exploratory approach, which similar to DISCO, applies SCA and can be used to decompose the original data set into a joint structure for the different data sources and

[1] Given two matrices of the same dimensions, the Hadamard product or element-wise product, is a binary operation that outputs a matrix of the same dimension as the operands containing the element-wise multiplication of their elements.

individual structures specific to each source. JIVE starts with a simultaneous component analysis of the concatenated matrices $X = [X_1 \dots X_K]$ and decomposes each omics data $X_k$ into the sum of three terms

$$X_k = C_k + D_k + E_k, \quad k = 1, \dots, K \qquad \text{(Eq. S32)}$$

where $C_k$ is the $I \times J_k$ matrix of $C$ related to the $k^{th}$ omics data, $D_k$ $(I \times J_k)$ is the individual pattern of $X_k$ and $E_k$ $(I \times J_k)$ gives the residual noise. Decomposition is achieved by an iterative procedure: at each step, the rank of $C$ is and $D_k$ $(k = 1, \dots, K)$ is estimated initially and then both matrices are computed by low-rank approximations of the concatenated matrix. In this step, the method attempts to minimize the sum of squared errors of the residuals $E_k$ under the given ranks, with the additional constraint $CD_k^T = 0$, to ensure that the orthogonality between the joint and individual structure is preserved. This is performed until convergence of residuals is succeeded. In contrast to DISCO, SCA in JIVE is performed only with the number of common components $(c)$ and not all of the them $(R)$. The resulting decomposition in scores and loadings though for two data matrices $X_1$ and $X_2$, is the same as for DISCO

$$\begin{aligned} X_1 &= C_1 + D_1 + E_1 = T_c P_{c_1}^T + T_{d_1} P_{d_1}^T + E_1 \\ X_2 &= C_2 + D_2 + E_2 = T_c P_{c_2}^T + T_{d_2} P_{d_2}^T + E_2 \end{aligned} \qquad \text{(Eq. S33)}$$

Again, the common scores $T_c$ are the same for both datasets.

*Two-way Orthogonal Partial Least Squares (O2PLS)*

Trygg and Wold[15] introduced the Two-way Orthogonal PLS (O2PLS), for decomposing the variation present in two data matrices ($X_1(I \times J_1)$ and $X_2(I \times J_2)$) into three parts, namely, the joint, the orthogonal and the noise part. The joint part represents the underlying mechanisms that are assumed to account for the variation that is common between the two matrices. The orthogonal part describes mechanisms that are assumed to account for the variation which is unique in $X_1$ and independent (i.e. orthogonal) to that in $X_2$.

In contrast to DISCO and JIVE, that initially perform an SCA on the concatenated matrices, O2PLS makes use of the covariance matrix $X_1^T X_2 (J_1 \times J_2)$ for an analysis of the common variation. First, the algorithm estimates the common components per dataset, and the distinctive components are estimated next orthogonal to the common part. After the common and distinctive variation is determined, they are removed from the data and the common scores are updated. With different common scores per dataset, the decomposition of O2PLS procedure for $X_1$ and $X_2$ in scores and loadings for $R$ components are similar to equations (S30) and (S33):

$$X_1 = C_1 + D_1 + E_1 = T_{c_1}W_{c_1}^T + T_{X_2\perp}P_{X_2\perp}^T + E_1$$
$$X_2 = C_2 + D_2 + E_2 = U_{c_2}Q_{c_2}^T + U_{X_1\perp}P_{X_1\perp}^T + E_2 \quad \text{(Eq. S34)}$$

where a linear relation exists between $T_{c_1}$ and $U_{c_2}$ and the score matrices are defined as

$$T_{c_1}(I \times c), T_{X_2\perp}(I \times d_1), U(I \times c), U_{X_2\perp}(I \times d_2) \quad \text{(Eq. S35)}$$

the loading matrices are defined as

$$W_{c_1}(J_1 \times c), C(J_2 \times c), P_{X_2\perp}(J_1 \times d_1), P_{X_1\perp}(J_2 \times d_2)$$

and the noise matrices are

$$E_1(I \times J_1),\ E_2(I \times J_2) \quad \text{(Eq. S36)}$$

where $c$ denotes the number of joint components, $c_1, c_2$ denote the joint structure for $X_1$ and $X_2$, $d_1$ the $X_1$-components that are orthogonal to $X_2$, and $d_2$ the $X_2$-components that are orthogonal to $X_1$.

*The simultaneous component analysis model*

SCA can be applied for exploring multiple data blocks together, and it is useful for estimating a few simultaneous components that represent most of the variability in the data. Consider $K = 2$ data matrices $X_1, \dots, X_k$, $(k = 1, \dots, K)$ that are to be integrated, with the same $I$ objects, samples, or experimental conditions and different $J_k$ variables in each matrix. The $K$ matrices can first be combined into a concatenated matrix as

$$X(I \times J) = [X_1(I \times J_1)| \dots |X_k(I \times J_k)],\quad k = 1, \dots, K \quad \text{(Eq. S37)}$$

with $J = J_1 + \dots + J_k$. Then, an SCA decomposition of the concatenated matrix $X$ into $R$ components is

$$X = TP^T + E \quad \text{(Eq. S38)}$$

where $T$ denotes the component score matrix of size $I \times R$, $P$ the matrix of the component loadings of size $J \times R$, and $E$ an $I \times J$ matrix of residuals. Then by separating the loadings $P$ into block-specific parts, each data block $X_k$ can be decomposed into

$$X_k = TP_k^T + E_k,\quad k = 1, \dots, K \quad \text{(Eq. S39)}$$

with $T^TT = \mathrm{I}$[2] and $T_1 = \dots = T_k = \dots = T_K = T\ (I \times R)$, $P_k$ the matrix of component loadings of size $J_k \times R$ and $E_k$ the residual error for the $k^{th}$ block. The SCA components represent underlying biological mechanism, however, they usually reflect a mix of common and

[2] The identity matrix, denoted by I is a square matrix with ones on the main diagonal and zeros elsewhere.

distinctive information, as the model is not able to separate common and distinct components. To further identify joint and specific variation in the composite data, each of the implemented integrative methods mentioned previously follows a specific methodology which is described in that same section.

The score matrix $T_k$ and loading matrix $P_k$ of the simultaneous component model with R components is estimated by the following optimization least squares function

$$\min_{T_k, P_k} \sum_k ||X_k - T_k P_k^T||^2 , \quad k = 1, \dots, K \qquad \text{(Eq. S40)}$$

with the constraint that $T_1 = \cdots = T_k = \cdots = T_K = T\ (I \times R)$, when the object mode is common between the data blocks. The optimal parameters for equation (S40) can be estimated by a singular value decomposition of the concatenated matrix $X$. SVD is a dimensionality reduction method that decomposes a matrix into the product of two unitary matrices[3] $(U, V)$ and a rectangular diagonal matrix with nonnegative real numbers on the diagonal (i.e. singular values) ranked in descending order $(S)$. Given the concatenated matrix $X$, the SVD decomposition is

$$X(I \times J) = USV^T \qquad \text{(Eq. S41)}$$

with $U\ (I \times I)$, $S\ (I \times J)$, and $V\ (J \times J)$. For a number of $R$ components and a common object mode (i.e. data matrices that share the same set of samples represented in the rows of the datasets), the score $T$ and the loading matrix $P$ for matrix $X(I \times J)$ are obtained as

$$\begin{cases} T = US \\ P = V^T \end{cases} \qquad \text{(Eq. S42)}$$

with $U\ (I \times R)$, $S\ (R \times R)$, and $V\ (R \times J)$.

[3] A unitary matrix $U$ is a squared matrix whose conjugate transpose is also its inverse, that is if $U^T U = UU^T =$ I, where I is the identity matrix.

#### Supplementary Note 2: Analysis for longitudinal gut microbiome profiles from children with Crohn's disease that undergo dietary treatment with EEN and healthy subjects

##### 2.2.1 Gut microbial community structure

When the samples from Crohn's disease patients and healthy individuals were grouped based on disease and healthy status (CD vs. Healthy), it could be seen that a few CD samples separated from the rest of CD points across the X axis (Fig. 2C, upper left-hand side). These samples were found to be associated with the patients that didn't achieve remission at point D of the treatment (Fig. S2), showing again that EEN can affect noticeably the community structure of the CD subjects.

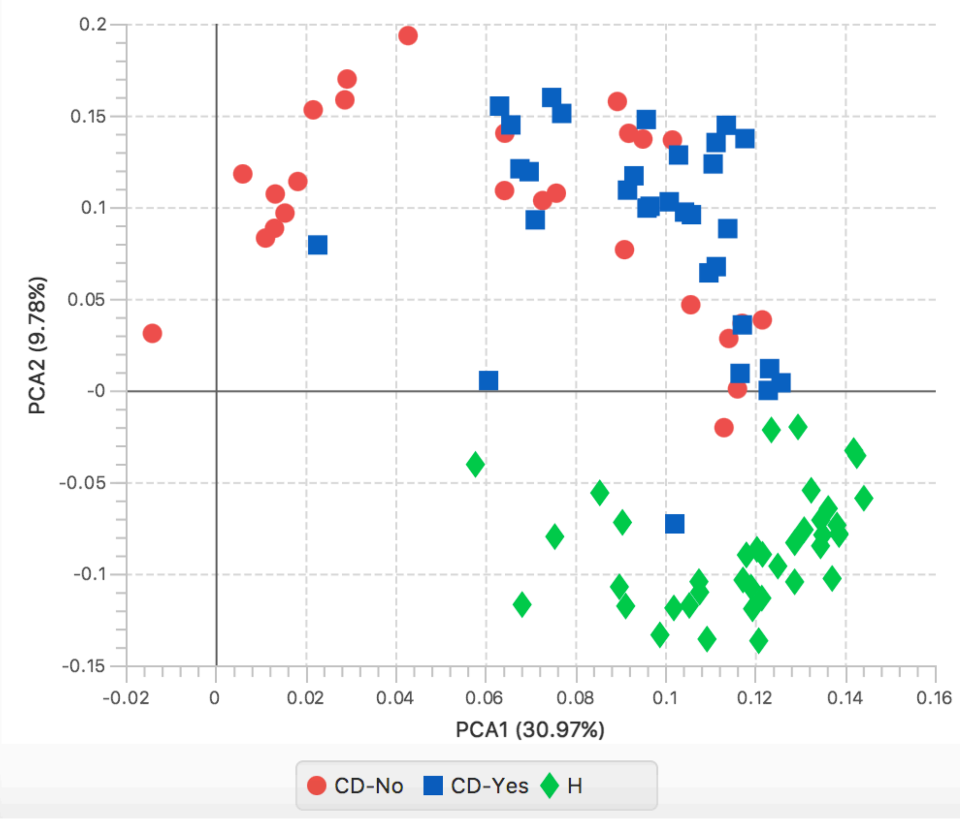


**Figure S2**: PCA analysis of shotgun metagenomics community structure of CD patients grouped according to whether they achieved remission at point D (End of diet) of EEN and healthy controls.

##### 2.2.2 Pielou's evenness

Evaluation of evenness (by Pielou's evenness) described a similar pattern for the CD samples during EEN, suggesting that the microbial communities of the patients became less even in abundance during the first phases (~30 days) of treatment and regress to pre-treatment levels after the end of EEN (Sample E) (Fig. S3). Compared to healthy groups, the bacterial community of CD patients was significantly less even at all samplings points (A:Start of EEN; $p=0.0001$, B:15d; $p<0.0001$, C:30d; $p<0.0001$, D:End of EEN; $p<0.0001$, E:Free Diet; $p=0.0034$).

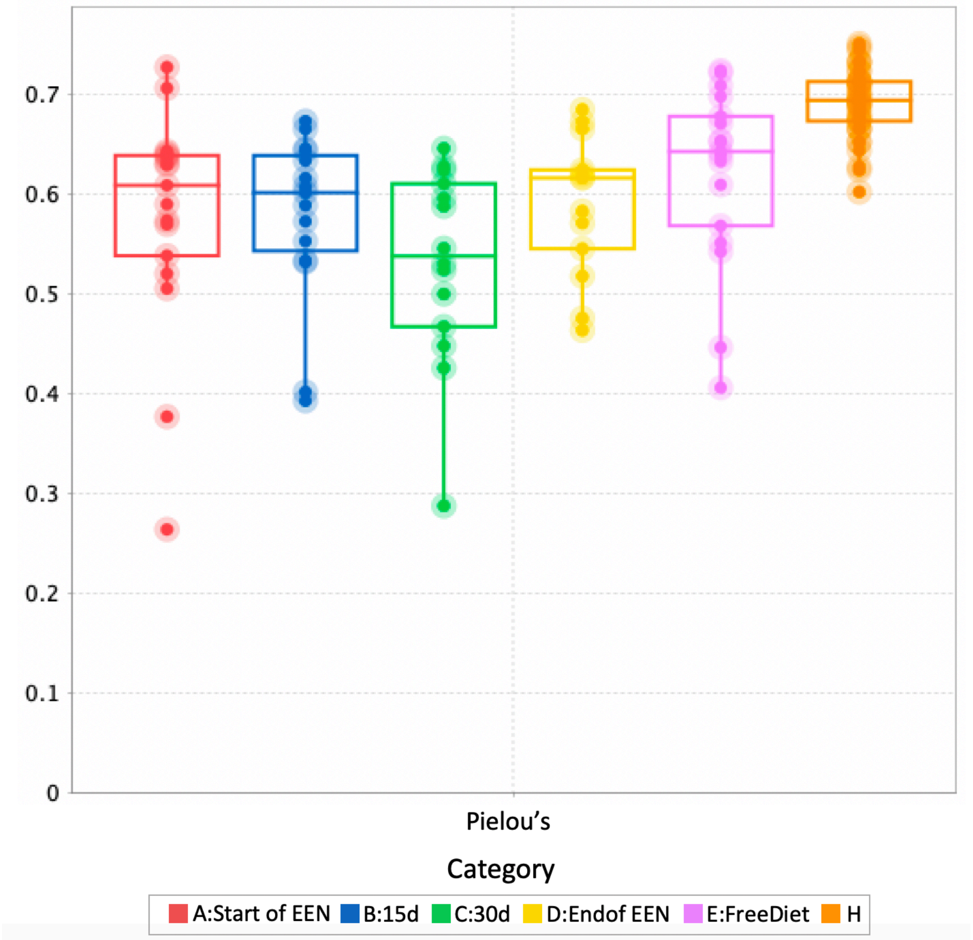


**Figure S3:** Pielou's evenness of CD patients during EEN and healthy controls. Pielou's evenness decreased over treatment and was lower in CD subjects compared to healthy controls. Changes reverted to pre-treatment levels when patients returned to their habitual diet.

##### 2.2.3 Correlations with faecal calprotectin

Fuzzy set ordination (FSO) analysis suggested an apparent relationship between the microbial community of the CD children with calprotectin levels, a marker of gut inflammation, at points D (end of EEN) and E (free diet) of EEN. Our findings suggested a significant correlation between the microbial community structure of the CD individuals at point D with calprotectin ($r=0.638$, $p=0.001$) and showed a distinct clustering associated with remission status (Fig. S4A). It was evident from the FSO plot, that patients who achieved remission were related to lower calprotectin levels compared with the patients who still have active disease following completion of EEN. A similar but less apparent pattern was noticed at point E, where calprotectin was also found to relate significantly with the community composition of the CD individuals ($r=0.574$, $p=0.001$) (Fig. S4B). This was not an unexpected outcome as CD patients regress to pre-treatment levels when they return to their habitual diet and as a result, changes induced by EEN become less apparent at that timepoint of treatment.

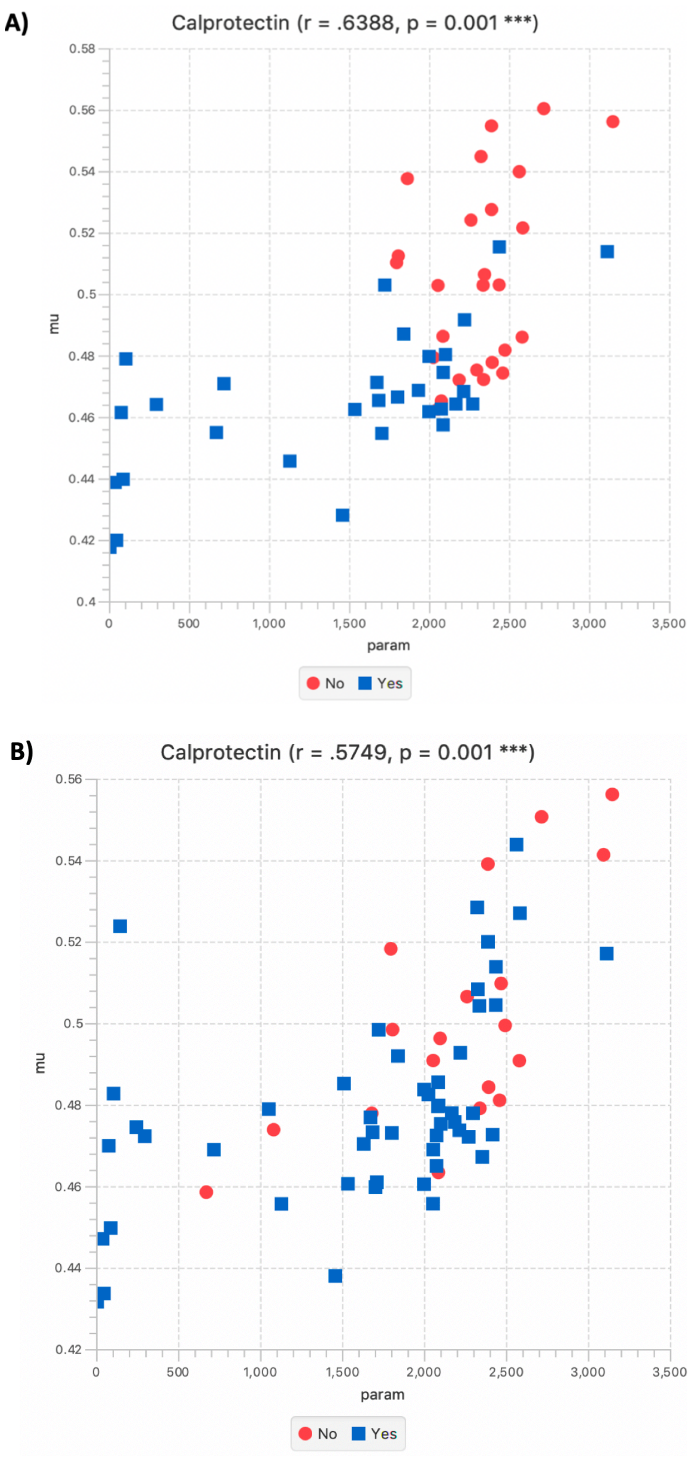


**Figure S4:** Fuzzy set ordination analysis demonstrating the relationship between calprotectin levels with the microbial community structure of CD subjects who were or not in remission at point D (end of EEN) (A) and at point E (free diet) of EEN treatment (B). The gut microbiota of CD patients clustered according to remission status and was significantly associated with calprotectin levels at both time points of treatment.

#### Supplementary Note 3: Analysis for gut microbiome profiles from individuals who are naturally and/or pathologically obese as compared to those who are lean

##### 2.3.1 Subject characteristics

When we compared the obese children according to pathology (healthy obese vs. hypothalamic obese) it could be seen that the standardized weight (Wt SDS), height (Ht SDS) and body mass index (BMI SDS) of healthy obese subjects was significantly higher than the hypothalamic obese individuals (BMI SDS, p=0.0158; Ht SDS, p=0.0009; Wt SDS, p=0.0004) (Data_Table_S3.xlsx). Height SDS was also significantly higher for the healthy lean controls compared to the hypothalamic lean participants (p=0.0004). However, BMI SDS of healthy lean individuals was significantly lower compared to the hypothalamic lean group (p<0.0001). When samples were grouped according to pathology, it was observed that pathology participants (hypothalamic lean and hypothalamic obese) were significantly shorter than the healthy individuals (healthy lean and healthy obese) (p<0.0001) suggesting a significant effect of pathology on height SDS. In addition, hypothalamic individuals had a significantly higher BMI SDS (p=0.0146) than the healthy groups.

##### 2.3.2 Gut microbial metabolites

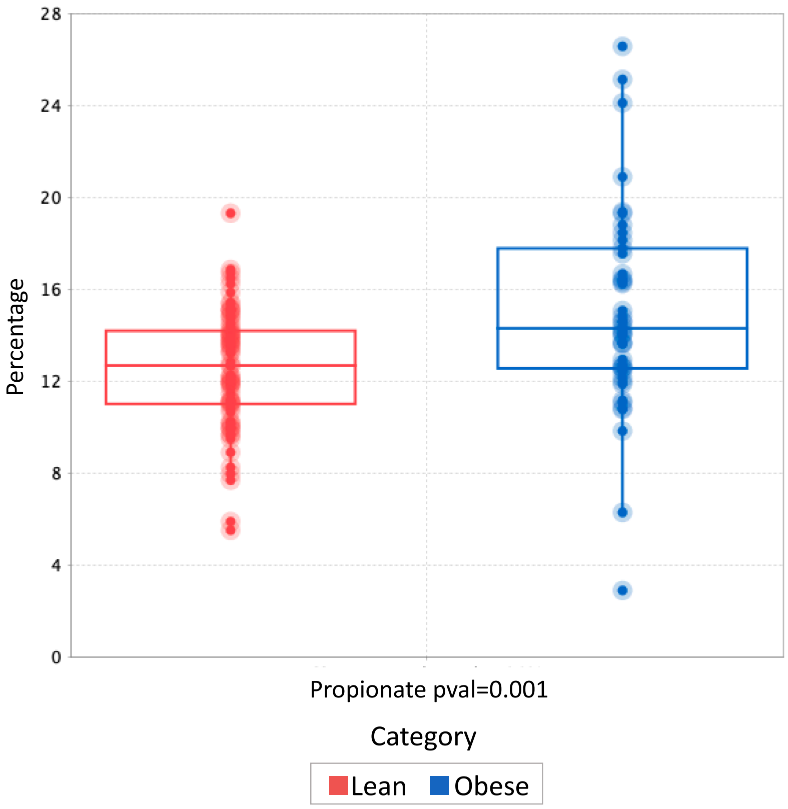


**Figure S5:** Box plots showing significant differences in the proportion of *propionate* according to lean and obese phenotype. Lean; healthy and hypothalamic lean, obese; healthy and hypothalamic obese.
